# A widespread family of phage-inducible chromosomal islands only steals bacteriophage tails to spread in nature

**DOI:** 10.1101/2022.09.08.507074

**Authors:** Nasser Alqurainy, Laura Miguel-Romero, Jorge Moura de Sousa, John Chen, Eduardo P.C. Rocha, Alfred Fillol-Salom, José R Penadés

## Abstract

Phage satellites interfere with helper phage packaging through the production of small-capsids, where only satellites can be packaged. So far, in all the analysed systems, the satellite-sized capsids are composed of phage proteins. Here we report the first demonstration that a family of phage-inducible chromosomal island (PICIs), a type of satellites, encodes all the proteins required for both the production of the small-sized capsids and the exclusive packaging of the PICIs into these capsids. Therefore, this new family, that we have named cf-PICIs (<u>c</u>apsid forming PICIs), only requires phage tails to generate infective PICI particles. Remarkably, the representative cf-PICI reproduces without cost for their helper phages, suggesting that the relationship between these elements is not parasitic but commensalistic. Finally, our phylogenomic studies indicate that cf-PICIs are present both in Gram-positive and Gram-negative bacteria and have evolved at least three times independently to spread widely into the satellite universe.

## Introduction

Mobile genetic elements (MGEs) are key players driving bacterial evolution and ecology (Koonin et al., 2019). Phage satellites are an important type of MGEs that couple their life cycle to that of the helper phages they parasitise, a strategy that ensures their promiscuous dissemination in nature at the expense of their helper phages. To date, the most studied phage satellites are P4 in *Enterobacterales* (Sousa and Rocha, 2022), the phage-inducible chromosomal islands (PICIs) in *Bacillales* and *Gammaproteobacteria* (Fillol-Salom et al., 2018; Martínez-Rubio et al., 2017), and the phage-inducible chromosomal islands-like elements (PLEs) in *Vibrio* spp. (O’Hara et al., 2017; Barth et al., 2019). Although the genetic organisation of these families of satellites is different, all these elements encode genes required for their regulation, replication and preferential package at the expense of their helper phages (Penadés and Christie, 2015; O’Hara et al., 2017; Christie and Dokland, 2012; Sousa and Rocha, 2022). Importantly, these elements do not encode the proteins required for the formation of their infective particles, and therefore, for their survival they must hijack the structural proteins and assembly processes of their helper phages (Ibarra-Chávez et al., 2021; Penadés and Christie, 2015).

Satellites have evolved different strategies to promote their preferential packaging into capsids composed of phage-encoded proteins. A strategy commonly used by satellite phages to interfere with phage packaging is the production of small capsids, which are commensurate to the size of the satellite’s genomes, but too small to accommodate the complete helper phage genome. Remarkably, although this strategy is conserved, satellites use different mechanisms to redirect phage capsid assembly, which represent a nice example of convergent evolution. While the satellite phage P4 expresses Sid (an external scaffolding protein) to redirect P2 capsid assembly (Agarwal et al., 1990), PICIs have evolved multiple mechanisms, depending on the phage they parasite (recently reviewed in (Fillol-Salom et al., 2020)). Thus, some staphylococcal pathogenicity islands (SaPIs), which are the prototypical members of the PICI family, express CpmB, which operates as a scaffolding protein, binding to the phage capsid protein, altering the curvature of the shells and redirecting its assembly and capsid size. This system also requires CpmA, whose predicted function is to remove the phage scaffolding protein (Christie and Dokland, 2012; Dearborn et al., 2017; Ubeda et al., 2007). Other SaPIs that use the *cos* system for packaging express Ccm, which has homology with the helper phage major capsid protein and redirects the assembly of the phage-encoded capsid protein into the production of SaPI-sized small capsids (Quiles-Puchalt et al., 2014; Carpena et al., 2016; Hawkins et al., 2021). PICIs from *Enterococcus faecalis* (EfCI583)(Matos et al., 2013), from *Pasteurella multocida* (PmCI172)(Fillol-Salom et al., 2018), and PLEs from *Vibrio cholerae* (Netter et al., 2021), have also the ability to produce satellite-sized capsids, although in these cases the proteins involved in this process have not been identified yet.

While capsid redirection severely reduces helper phage reproduction, it does not increase *per se* the packaging of the satellite phages into the small capsids. To solve this problem, satellites have evolved an arsenal of complementary and sophisticated strategies that ensure their preferential packaging into the virions, facilitating their promiscuous transfer in nature. Phage packaging starts by the recognition by the phage-encoded small terminase (TerS) of the *cos* or *pac* sites in the phage genomes. To promote their packaging, some satellites - such as P4 and some SaPIs - carry the cognate helper phage *cos* site in their genomes (Quiles-Puchalt et al., 2014; Ziermann and Calendar, 1990). Other SaPIs encode a homologue of the phage TerS called TerSS (Ubeda et al., 2007), which recognises the SaPI *pac* sequence (Chen et al., 2015). To help TerSS function, these SaPIs also encode Ppi (<u>p</u>hage <u>p</u>ackaging interference), which binds to the phage TerS, blocking its function (Ram et al., 2012, 2014). Finally, some PICIs present in Proteobacteria encode Rpp (redirecting <u>p</u>hage <u>p</u>ackaging), which binds to the phage TerS. The Rpp-TerS complex recognises the PICI *cos* site (promoting PICI packaging), but not the phage *cos* site, thus blocking phage reproduction (Fillol-Salom et al., 2019).

While diverse, all the mechanisms of piracy described so far require that helper phages express all the proteins necessary to generate not just the phage but also the PICI infective particles. We report here the discovery of a widespread family of PICIs, that we have named cf-PICIs (<u>c</u>apsid forming PICIs), that encode not just all the proteins required for the production of PICI-sized capsids but also for the exclusive packaging of the cf-PICIs into the small-sized capsids. Therefore, and contrary to what is seen with other satellites, cf-PICIs only require phage tails to spread in nature. They represent a new strategy of molecular piracy.

## Results

### EcCIEDL933 is the prototypical member of the cf-PICI family

During the description of PICI elements in *Proteobacteria* (Fillol-Salom et al., 2018), one intriguing element with an unusual genetic organisation, present in the *E. coli* strain EDL933, raised our curiosity. This element, with a size of 15.4 kb, carried the *int*, *alp*A and *rep* genes, identical to those present in the prototypical *E. coli* EcCICFT073 and EcCIED1a PICIs (Fig. 1A). However, in addition to these conserved PICI genes, this element seemed to encode all the proteins required for the production of functional capsids (major capsid, head maturation protease, portal, head-tail connectors), and for the packaging of the PICI DNA into these capsids (HNH and small and large terminase proteins) (Fig. 1A). Interestingly, this element did not encode genes involved in the production of phage tails.

**Figure 1.**
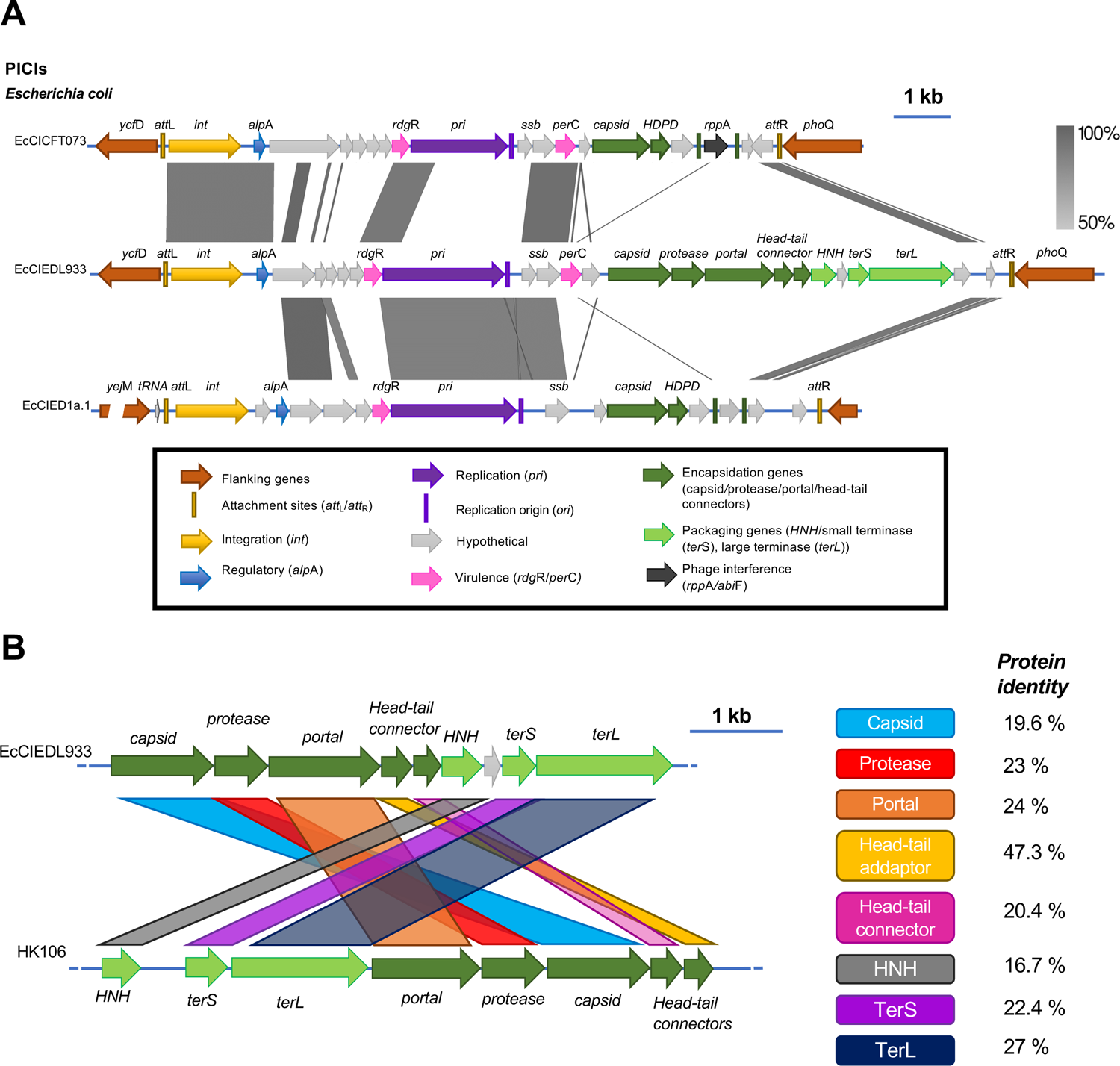
Comparation of the different PICI lineages. (A) Comparative maps of the classical *E. coli* PICIs EcCICFT073 and EcCIED1a with EcCIED933. Genomes are aligned according to the prophage convention, with the integrase genes (*int*) present at the left end of the islands. Genes are colored according to their sequence and function (see box for more details). Using Easyfig alignment, vertical blocks between PICI sequences indicate regions that share similarity based on BLASTn (grey scale). (B) Comparison of the packaging modules present in either EcCIEDL933 or in helper phage HK106, at protein level, using BLASTp. Identities are shown for each encoded protein.

Based on this peculiar organisation, our initial thoughts were that this element could represent a hybrid between a PICI and a phage. Note that the aforementioned DNA packaging genes present in this putative hybrid element showed homology with phage genes (Fig. 1B). Yet, sequence similarities between the genes present in this new element and those present in phages were generally low and restricted to a few genes. To assess the similarity in gene repertoires between the element present in strain EDL933, hereinafter named EcCIEDL933, and the genomes of typical phages, we computed their weighted gene repertoire relatedness, which we have used previously to class phages in families (see Methods)(Sousa et al., 2021). Our analysis of 3725 complete non-redundant phage genomes retrieved from RefSeq shows that the genome of EcClEDL933 is very different from them. Indeed, 86% of the phage genomes lack homologs in EcClEDL933, with the remaining 14% (519 phage genomes) having, at most, a wGRR of 0.09 (Fig S1). These cases represent phage-encoded homologs to some of the proteins encoded by EcClEDL933, and include, besides the aforementioned capsid and terminase genes, homologs to the other capsid formation (portal, head-tail connectors) or packaging (HNH) genes. Importantly, no single phage genome has more than six homologous proteins with EcClEDL933 and the average number of homologs, among phages with at least one homolog, is only 1.8 (Fig. S1D). Most of these rare homologs have significant, but low sequence similarity (<40%, e-value<1e-04; Fig. S1C and S1D). On the other hand, our KEGG orthology analyses showed that genes homologous to the ones present in EcClEDL933 are frequently found in genomic regions that could correspond to similar elements in *E. coli* and other related species, including *Salmonella enterica, Shigella flexneri, Klebsiella pneumoniae, Pluralibacter gergoviae* or *Yersinia aleksiciae* (Table S1). Therefore, these results suggested that EcCIEDL933 could represent a new family of elements, distinct from both phages and previously described PICIs. In support of this idea, members of this new family integrate into the *E. coli* bacterial chromosome using six different *att*B sites. Of these, two were exclusively used by the members of this new lineage, while the other four were also used by members of the previously identified *E. coli* PICIs (Table S2 and Fig. S2).

The previous results suggested that the mechanism of induction of this new lineage could be identical to that described for the classical PICIs present in *E. coli*. Once induced, the genes involved in DNA packaging could either promote the packaging of the cognate element and/or could be used by the island to interfere with helper phage reproduction. To clearly demonstrate that EcCIEDL933 is induced as the classical PICI elements, we searched for helper phages able to induce and mobilise EcCIEDL933. To facilitate these studies, we inserted a *cat* marker into the element present in the clinical EDL933 strain, which also carries 17 prophages (Perna et al., 2001), and induced the resident prophages with mitomycin C (MC), expecting that one of the resident prophages was able to mobilise EcCIEDL933 *cat* into the non-lysogen 594 *E. coli* strain, used in this experiment as recipient. This was the case, although just a few transductants were obtained (Table 1), suggesting that none of the resident prophages present in the EDL933 strain are the helper phage for this island. Note that in presence of a helper phage, PICI transfer is extremely high (Humphrey et al., 2021). Thus, it is likely that neither of the 17 prophages induces EcCIEDL933.

**Table 1.**
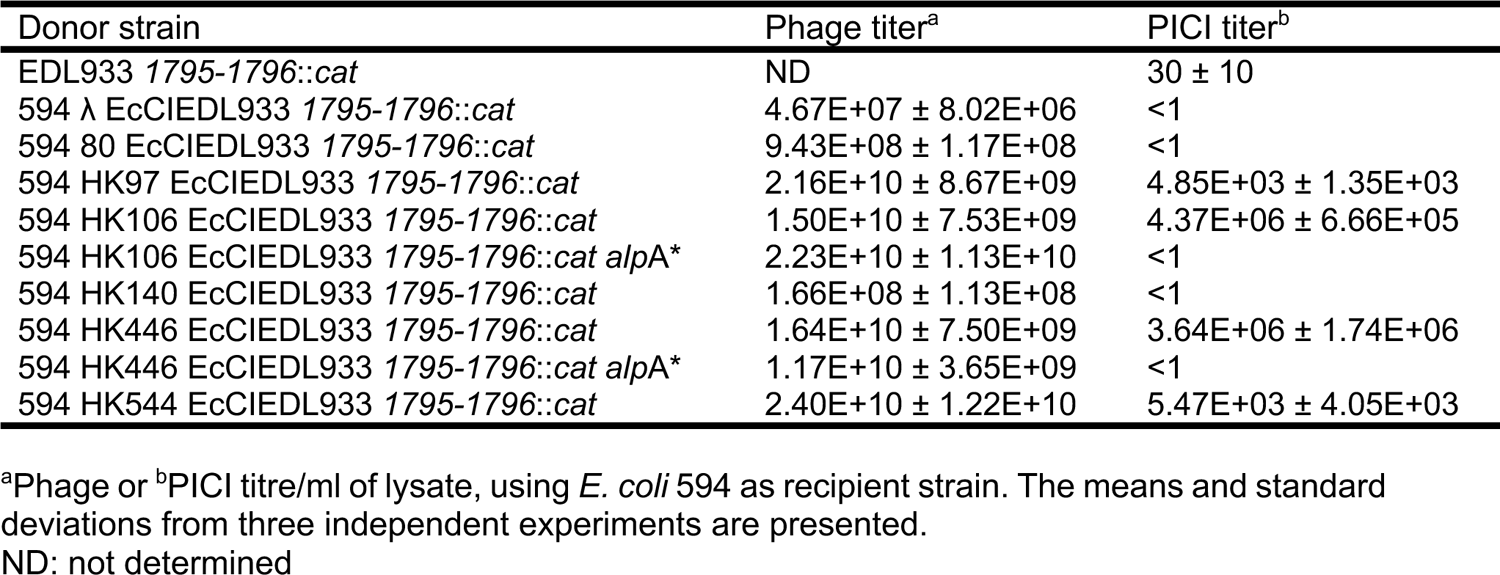
EcCIEDL933 transfer by different *E. coli* phages.

To search for one, we lysogenised the *E. coli* 594 strain carrying EcCIEDL933 *cat*-positive with our collection of *E. coli* phages and tested the transfer of the PICI element after MC induction of the different lysogenic strains. Two phages, HK106 and HK446, mobilised the island at high frequencies (Table 1), suggesting they were able to induce the EcCIEDL933 cycle. This was confirmed with the screening lysate analysis of the DNA samples obtained after induction of the different lysogenic strains carrying EcCIEDL933. As shown in Fig. 2, only the strains lysogenic for either HK106 or HK446 showed the classical pattern of PICI induction: a high sized DNA band corresponding either to the PICI replicating concatemeric form or to the PICI/phage DNA packaged into the phage-sized capsids, and a small band corresponding to the PICI DNA packaged into small PICI-sized capsids.

**Figure 2.**
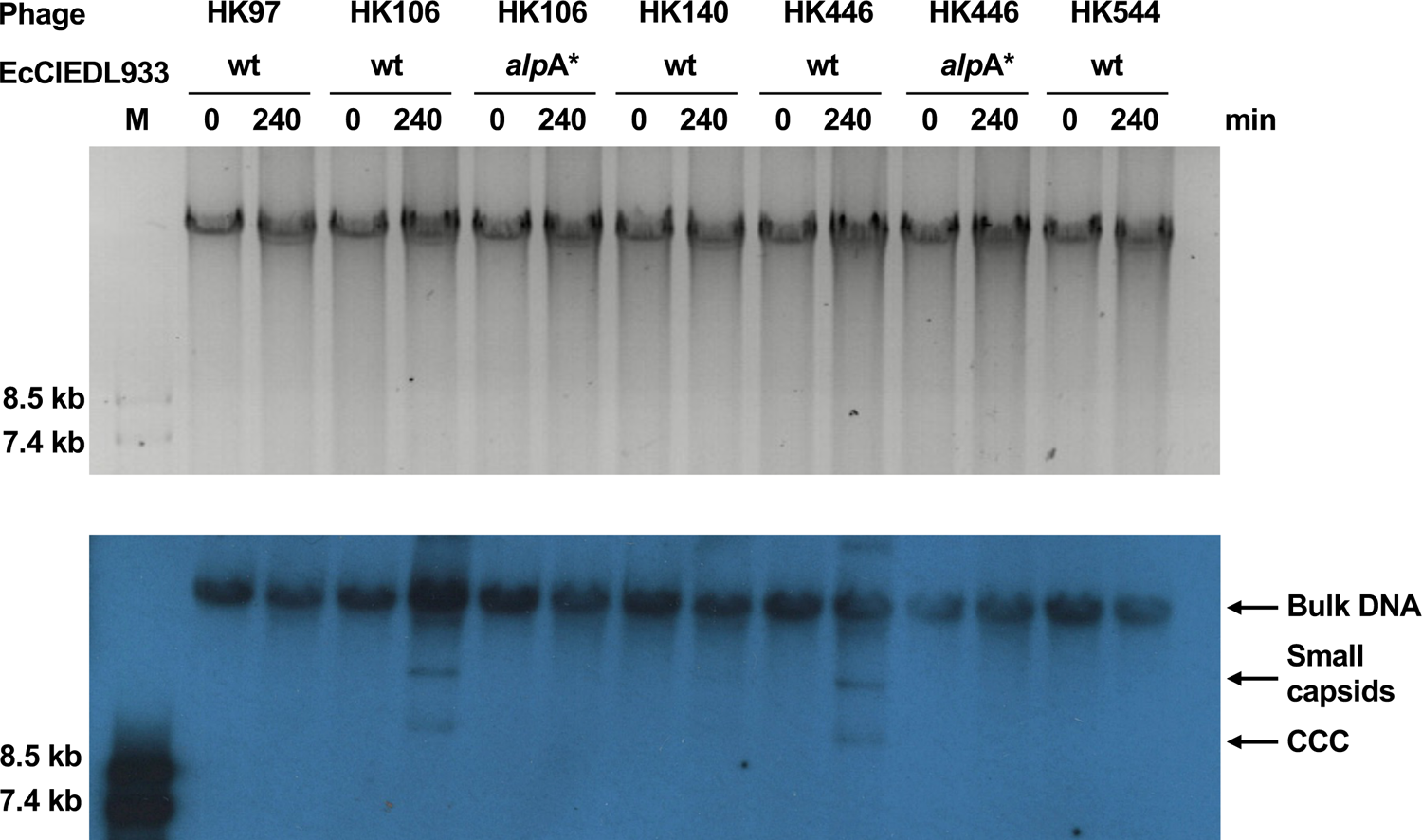
Phage induction of EcCIEDL933. Different lysogenic strains carrying EcCIEDL933 (wt or mutant in *alp*A (*alp*A*) were MC (2 μg/ml) induced, samples were taken at the indicated time points (min) and processed to prepare minilysates, which were separated on a 0.7% agarose gel (upper panel), and Southern blotted (lower panel) with an EcCIEDL933 probe. M: Southern blot molecular marker (DNA molecular weight marker VII; Roche).

Finally, we tested the transfer of the EcCIEDL933 *alp*A mutant. Note that in Gram-negative bacteria PICI induction and transfer requires the expression of the PICI-encoded *alp*A gene, a process that is activated by the helper phage (Fillol-Salom et al., 2018). As expected for a PICI-like element, mutation of *alp*A eliminated both helper-phage induction (Fig. 2) and transfer of this element (Table 1).

### EcCIEDL933 produces PICI-sized capsids

Having established that this new family of satellites are induced like the classical PICIs, we wanted to know whether the EcCIEDL933 genes with homology to the phage genes were required for capsid formation and EcCIEDL933 packaging and transfer. The presence of the PICI-sized small band in the EcCIEDL933 screening lysate suggested that this island had the ability to produce PICI-sized capsids after induction. To test this, we analysed by electron microscopy (EM) the infective particles present in the lysate obtained after induction of the lysogenic strain for helper HK106, in presence or absence of EcCIEDL933. In the absence of EcCIEDL933, only phage particles were observed, which had the characteristic size and shape of a *Siphoviridae* phage with a hexagonal capsid of approximately 60 nm length and 57 nm width, with non-contractile tails which are approximately 160 nm long and 10 nm wide (Fig. 3A). By contrast, in the presence of EcCIEDL933 two different types of infective particles were observed: in addition to the phage-sized ones, we were able to detect PICI-sized capsids, which had a hexagonal capsid of 46 nm length and 42 nm wide, connected to the phage non-contractile tail (Fig. 3A).

**Figure 3.**
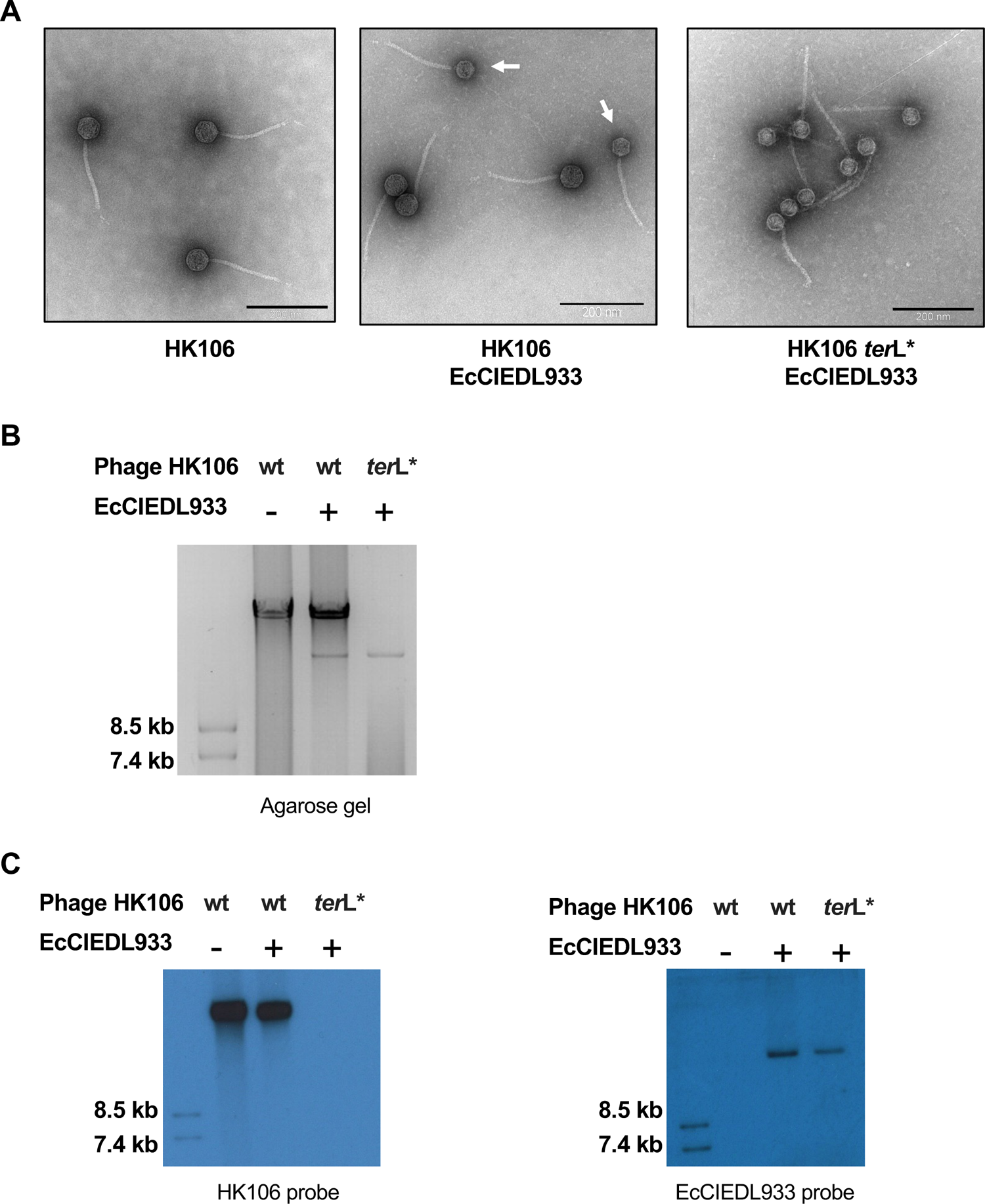
EcCIEDL933 produces small capsids. (A) Electron microscopy analyses of different MC-induced HK106 lysogenic strains (wt or *ter*L mutant (*ter*L*) in presence or absence of EcCIEDL933. Different fields are shown, containing HK106 phage particles (left), EcCIEDL933 particles (right) or both (middle; PICI particles highlighted with white arrows). EcCIEDL933 particles have smaller heads. Scale bars are 200 nm. (B) Lysogenic strains carrying the wt or the *ter*L mutant (*ter*L*) HK106 prophage, in presence or absence of EcCIEDL933, were MC-induced, the DNA was extracted from the purified infective particles and resolved on a 0.7% agarose gel. (C) Southern blot of the purified DNA shown in panel (B) using either phage HK106 or EcCIEDL933 specific probes. First line contains the Southern blot molecular marker (DNA molecular weight marker VII; Roche).

Since the PICI genome is 1/3 of the size of the phage genome, it is obvious that the complete phage genome cannot be packaged into the PICI-sized particles. However, some PICIs can package 3 copies of their genomes in phage-sized capsids (Carpena et al., 2016; Ubeda et al., 2007). To test whether this was also the case for EcCIEDL933, the packaged DNA present in the lysate containing both the phage and the PICI infective particles was purified, separated on agarose gels, and analysed by Southern blot, using specific phage- or PICI probes. In support of the EM images, two types of bands were observed, corresponding to the size of the phage and EcCIEDL933 genomes, respectively (Fig. 3B). Importantly, the Southern blot analyses revealed that the packaging of the different elements is size specific and mutually exclusive, with phages and PICIs being packaged uniquely in large- and small-sized capsids, respectively (Fig. 3C).

### Characterisation of the EcCIEDL933 packaging module

The previous results suggested that the different sized capsids had different origins: the phage would be responsible for the production of the large capsids and the phage tails, while the small capsids would be produced by the PICI. In support of this idea, comparison of the phage and PICI packaging modules reveals that they have different genetic organisations and low protein identities (Fig. 1B). To test this idea, we individually mutated each of the genes belonging to the putative packaging module of EcCIEDL933 (Fig. 4A) and tested the transfer of the different EcCIEDL933 mutants after induction of the helper prophage HK106. Remarkably, while the replication of the different mutants was unaffected (Fig. S3), their transfer was completely abolished in all cases except in the mutant for the hypothetical gene 1792, whose transfer was severely affected (Fig. 4B). To confirm that the observed phenotypes were consequence of the mutations, complementation of the mutants restored PICI titres (Fig. S4C). Interestingly, and in support of our idea, in all these cases the transfer of the helper phage was unaffected, confirming that the PICI encoded proteins have no role in the packaging of the helper phage (Fig. 4C). Importantly, the Southern blot analyses of the purified infective particles obtained after induction of the different EcCIEDL933 mutant strains uniquely revealed the presence of the phage-sized band, but not the PICI-sized ones, confirming that the small capsids had a PICI origin (Fig. S5C). Next, we performed the same experiment, but now mutating the phage genes encoding the homolog proteins to those encoded by the PICI. In this case the mutations eliminated phage but not PICI transfer (Fig. 4E and 4F). Phage mutants were complemented *in trans*, confirming that the phenotypes were due to the mutations (Fig. S4D). Moreover, and in agreement with the idea that the large and small capsids had different origins, only PICI-sized bands were observed in the Southern blot analyses of the purified infective particles obtained after induction of the different phage HK106 mutant strains (Fig. S5D). This was corroborated with the analysis of the lysate obtained after induction of the strain carrying the HK106 capsid (*gp05*) mutation and EcCIEDL933, which only contained PICI-sized particles (Fig. 3).

**Figure 4.**
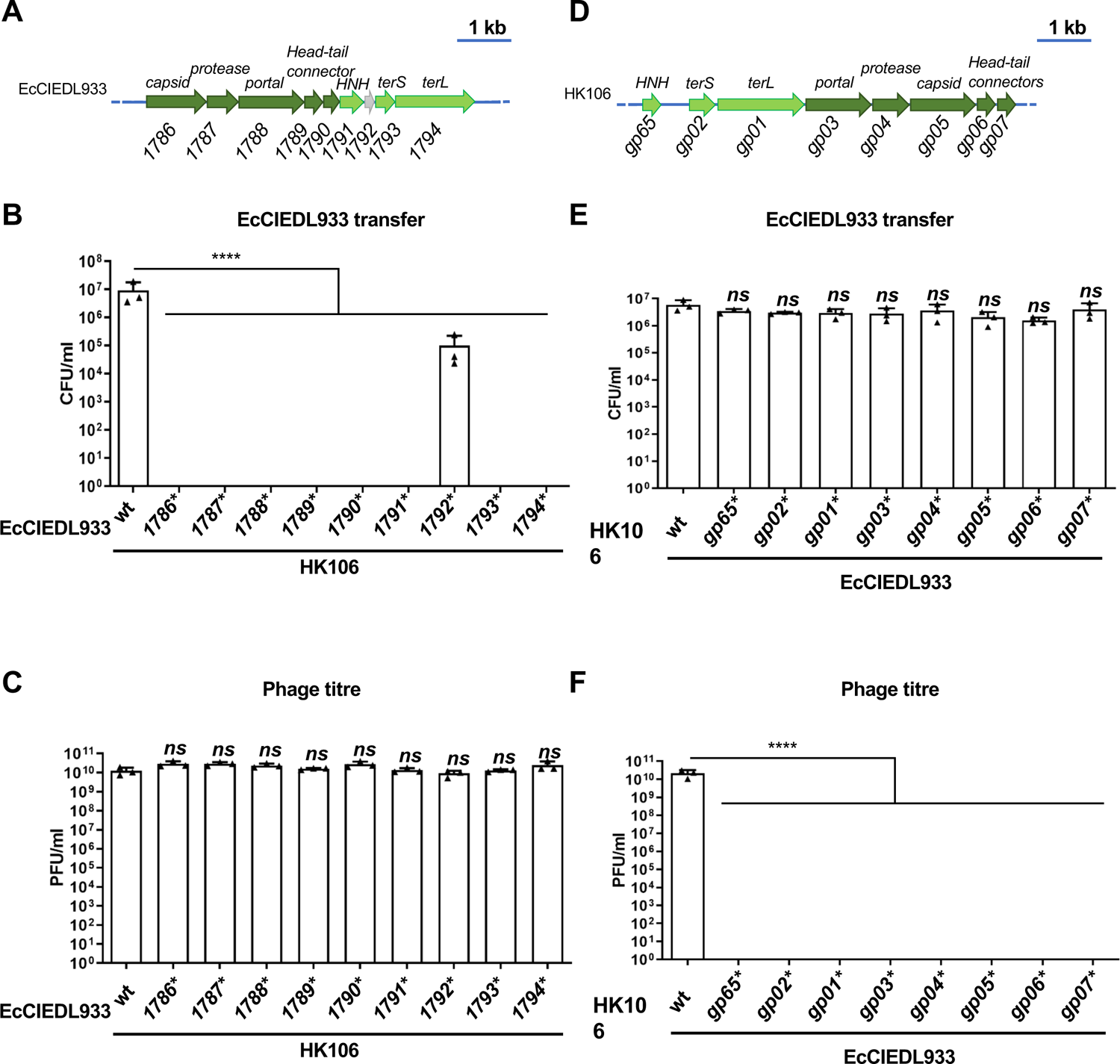
Effect of different PICI and phage gene mutations on EcCIEDL933 and HK106 transfers. Genetic maps of the EcCIEDL933. **(A)** or the phage HK106 **(D)** packaging modules. Lysogenic strains for phage HK106, carrying different EcCIEDL933 versions (wt or carrying mutations in the indicated genes), were MC (2 µg/ml) induced and the PICI **(B)** or phage **(C)** transfers were determined using *E. coli* 594 as recipient strain. Lysogenic strains for HK106, carrying the wt or the different mutant prophages, were MC (2 µg/ml) induced in the presence of EcCIEDL933 and the transfer of the island **(E)** or the phage titres **(F)** were determined using *E. coli* 594 as recipient strain. The means of the colony forming units (CFUs) or phage forming units (PFUs) and SD of three independent experiments are presented (n=3). A one-way ANOVA with Dunnett’s multiple comparisons test was performed to compare mean differences between wt and individually point-mutations. Adjusted p values were as follows: *ns*>0.05; *p≤0.05; **p≤0.01; ***p≤0.001; ****p≤0.0001.

Our previous results demonstrated that EcCIEDL933 encodes a functional packaging module responsible for both the production of the PICI-sized capsids and for the specific and exclusive packaging of the island into these particles. Therefore, we propose here that EcCIED933 is the prototypical member of a new family of PICI satellites, that we have named capsid forming PICIs (cf-PICIs), with the ability to produce all the proteins required for the specific packaging of these elements into cf-PICI encoded capsids. These results also suggested that in terms of producing infective particles, EcCIEDL933 only requires the phage tails to complete the formation of the PICI infective particles. To confirm this hypothesis, we mutated two of the phage genes encoding important tail components: the tape-measure (*gp14*) and major tail (*gp10*) proteins. The strains carrying the mutant phages and EcCIEDL933 were MC induced, and the phage and PICI titres quantified. As expected, these mutants abolished both phage and PICI transfer (Fig. 5A). Next, to confirm that EcCIEDL933 only relies on the phage tail proteins to produce PICI infectious particles, we analysed by EM the lysate obtained after the induction of a strain containing the PICI and a helper phage carrying mutations in the tape-measure (*gp14*) and the capsid (*gp05*) genes. Note that in this strain the phage does not produce large capsids nor tails. In support of our hypothesis, only PICI-sized capsids were observed in the lysate (Fig. 5B).

**Figure 5.**
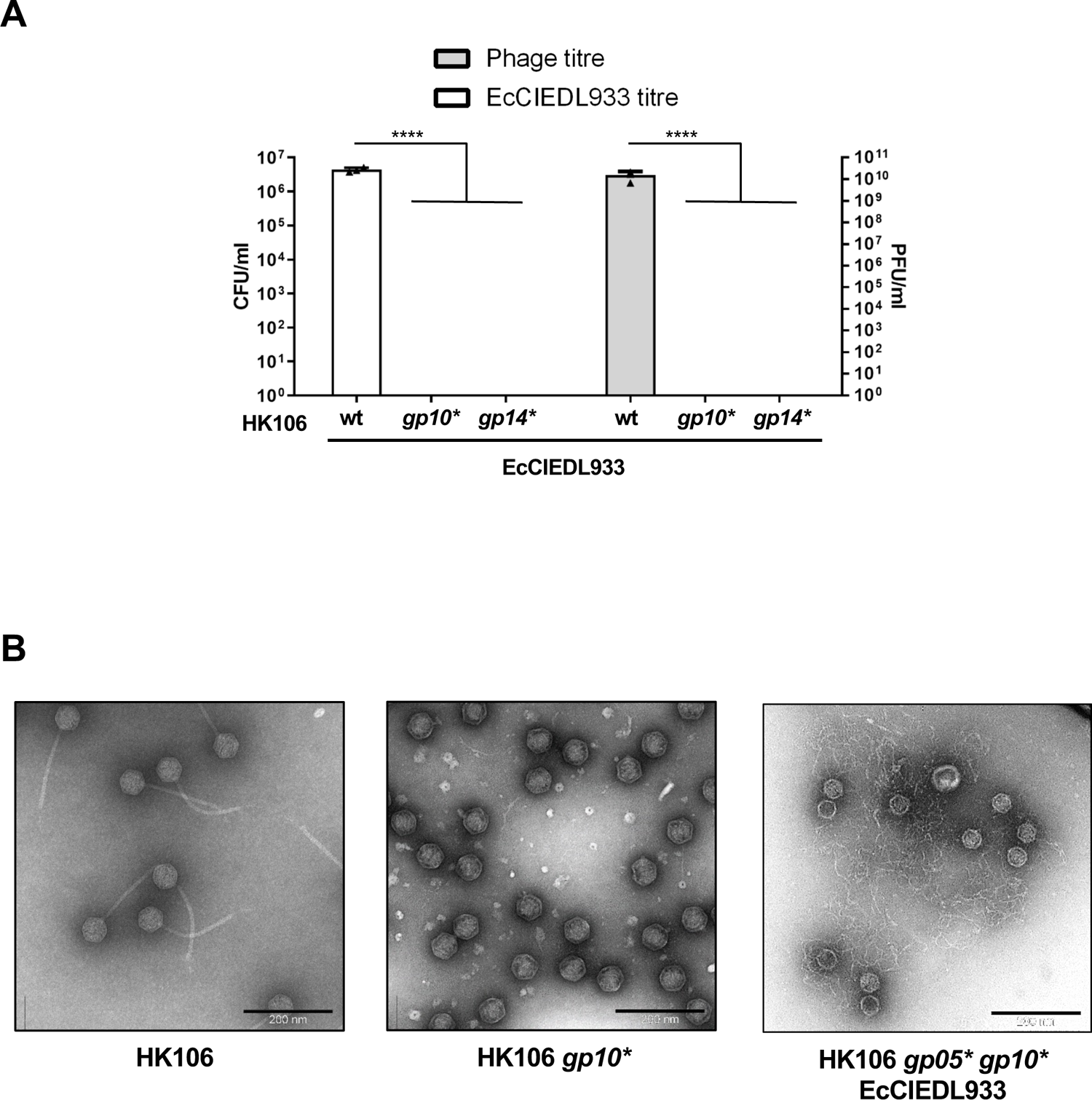
EcCIEDL933 requires phage tails to generate infective particles. (A) Lysogenic strains for phage HK106 (wt, Δ*gp10* or Δ*gp14*) carrying EcCIEDL933 were MC induced (2 µg/ml) and the phage and PICI titres were determined using *E. coli* 594 as recipient strain. Note that *gp10* and *gp14* encode major tail and tape-measure proteins, respectively, required for phage tail formation. The means of the colony forming units (CFUs) or phage forming units (PFU) and SD of three independent experiments are presented (n=3). A one-way ANOVA with Dunnett’s multiple comparisons test was performed to compare mean differences between wt and individually point-mutations. Adjusted p values were as follows: *ns*>0.05; *p≤0.05; **p≤0.01; ***p≤0.001; ****p≤0.0001. (B) Electron microscopy analysis of different MC-induced HK106 lysogenic strains (wt and mutants) in presence or absence of EcCIEDL933. Different fields are shown, containing only HK106 phage particles, HK106-sized capsid particles or EcCIEDL933-sized capsid particles. EcCIEDL933 particles have smaller heads. Scale bars are 200 nm.

### EcCIEDL933 does not block helper phage reproduction

One conserved characteristic of the satellites is their ability to severely interfere with their helper phage reproduction because they hijack its components (Fillol-Salom et al., 2020) (O’Hara et al., 2017). To test whether EcCIEDL933 was also able to block the reproduction of its helper phages, we quantified the phages obtained after induction of the lysogenic strains for HK106 or HK446, the EcCIEDL933 helper phages, in presence or absence of the island. Importantly, phages HK106 or HK446 were unaffected by the presence of EcCIEDL933 (Table S3). To confirm this, we infected strains 594 or JP22295 (a derivative 594 strain carrying EcCIEDL933) with phages HK106 or HK446, and compared their efficiency of plating. With the classical PICIs, in presence of the islands, both the number and the size of the phage plaques are severely reduced, only being able to plaque those mutant phages that are unable to induce the islands (Frígols et al., 2015). In this case, however, the scenario was completely different, and as occurred after prophage induction, the presence of the island did not affect the capacity of the helper phages to reproduce normally in the recipient strain carrying the island, compared to the EcCIEDL933-negative strain (Fig. S6). These results suggest that EcCIEDL933 is not a phage parasite, but rather a commensal

### cf-PICIs are widespread in nature

We next analysed whether this new family of satellite phages is widespread in nature. We made a curated search across the GenBank database for similar cf-PICI elements among thirteen species of *Proteobacteria*, especially in members of the *Enterobacteriaeae, Pasteurellaceae* and *Orbaceae*, where PICIs are frequent (Fillol-Salom et al., 2018). We searched for elements carrying *int* or *alp*A genes identical to those present in the canonical PICIs, but encoding a packaging module similar to that present in the cf-PICI EcCIEDL933. Our preliminary analysis revealed that other cf-PICIs are frequent in *Proteobacteria* genomes (Fig. S2; Table S2). Importantly, all the elements identified encode all, or most, of the components present in EcCIEDL933, with a similar conserved genetic organisation (Fig. 1 and S2). Their genes encoding the packaging module present sequence similarity with those from EcCIEDL933 ranging from 50% to 90%, suggesting that they are indeed part of a single family of satellites.

Using the same strategy that allowed us the identification of the cf-PICIs in *Proteobacteria*, we next interrogated the genomes of Firmicutes, the other phylum with known PICIs for the presence of these elements. Importantly, our analysis was able to identify cf-PICIs in different genera of Firmicutes, including *Lactococcus, Enterococcus, Bacillus*, *Clostridium* and *Staphylococcus*. While the cf-PICIs in *Proteobacteria* showed a conserved genetic organisation, with the morphogenetic genes (capsid-protease-portal-head connectors) first, followed by the ones involved in DNA recognition and packaging (*hnh*-*ter*S-*ter*L), two distinct groups were observed in the cf-PICIs from Firmicutes. One group was composed by the elements present in *Bacillus*, *Clostridium* and *Staphylococcus*, while the second group was composed by the elements present in *Lactococcus* and *Enterococcus* (Fig. S7; Table S4). While both groups had similar gene content, the organisation/localisation of these genes within the packaging operon was different. The genes encoding the portal, protease and capsid proteins of the first group were present at the beginning of the operon, while the genes encoding HNH, TerS and TerL start the operon of the second group. This suggests that cf-PICI may have different origins in these two phyla.

### cf-PICIs arose three times independently

To detail the evolutionary history of cf-PICI, we searched specifically for homologs of their capsid and large terminase proteins in the database of complete phage genomes used in our initial analysis. In each case (capsid or TerL), we collected the phage hits with highly significant e-values (<1e^-10^) (see Methods). This allowed us to put together a list of homologs for the capsid (113 phage + 61 cf-PICI) and TerL (420 phage + 60 cf-PICI) proteins present in the cf-PICIs or in the phages. We then built the phylogenetic trees of the two types of proteins. The conclusions were similar for the analyses of the capsid and the terminase proteins (Fig. 6 and S8). We observed that cf-PICIs regrouped on three clades that are clearly separated from each other by many phage genes. These three clades correspond to the three groups distinguished in the previous section in terms of genetic organisation. The separations between the three groups are robust to phylogenetic reconstruction uncertainty, as attested by high confidence bootstrap values at the basis of each clade. This suggests the independent emergence of the three groups of cf-PICIs.

**Figure 6.**
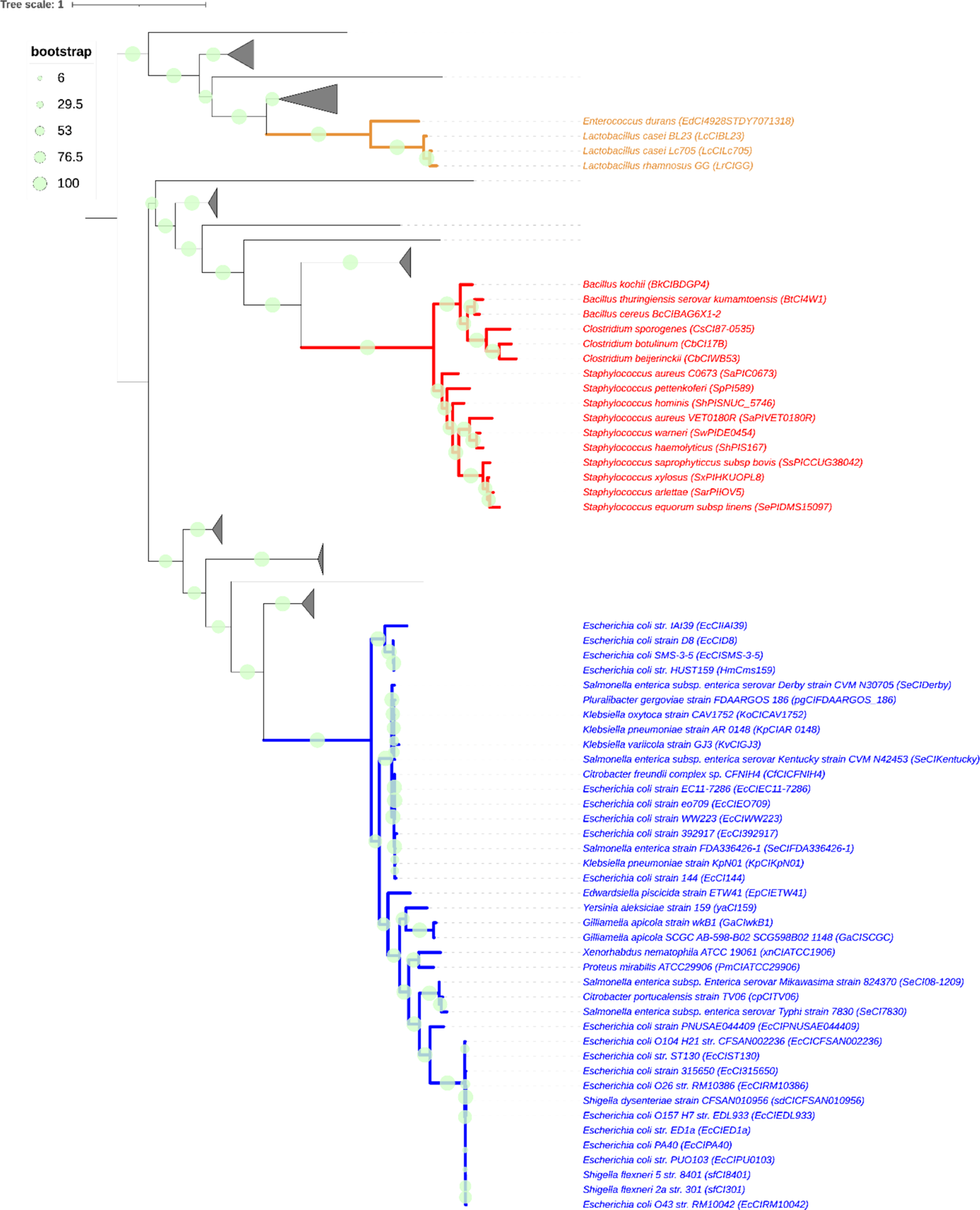
Capsid of cf-PICI form three distinct phylogentic groups and are evolutionarily separated from phage homologs. Phylogenetic trees inferred from the alignment of 61 capsid homologs from cf-PICI and the best 113 phage capsid homologs. Phages branches are collapsed and shown as triangles. The different colors on the branches indicate the three different clades, formed by either *Gammaproteobacteria* (in blue), *Lactobacillus* (in orange) and a broader set of *Firmicutes* (in red). Bootstrap values for the main branches are indicated as grey circles. The tree was visualized and edited in iTOL (Letunic et al., 2021). Phylogenetic tree resulting from the alignment of Capsid proteins from 61 cf-PICIs and 3725 phage.

The analysis of each of the cf-PICI clade shows a clear separation between the satellites and the phage homologs, i.e. the clades only include cf-PICI components even if we deliberately selected the phage homologs that were most similar to the cf-PICI. Furthermore, the phages closest to each clade of cf-PICIs tend to be the same in the TerL and capsid tree. The consistent phylogeny for these components suggests strong genetic linkage between the two functions. This further confirms that cf-PICIs are clearly distinguishable from the phages in the database and suggests a low frequency of genetic exchanges of these key components between cf-PICIs and phages.

Within each clade of cf-PICI one identifies elements present in the genomes of multiple bacterial genuses. In the EcCIEDL933-containing clade we could identify bacteria from very different Gammaproteobacteria, from *E. coli* to *Yersinia* or *Xenorhabdus*. Similarly, the clade containing the *S. aureus* element includes bacteria from *Bacillus* and *Clostridium*, which are very distantly related. This shows that either these elements have an exceptionally broad host range upon transfer through virions, or that they are very ancient and have diversified within distinct bacterial taxonomic groups. The analysis of the distances in the phylogenetic trees shows very long branches between the cf-PICI and the closest phages even in the highly conserved protein TerL (Fig. 6), which is also suggestive of an old origin for cf-PICIs. Hence, the cf-PICIs are neither recently derived from phages nor the result of recent prophage genetic degradation. Instead, they constitute deep lineages of elements that, compared to the other previously described PICI, have more proteins that are homologous to those of phages.

## Discussion

Satellites are MGEs whose life cycle depends on a helper virus but lack extensive nucleotide sequence homology between them. So far, what is the origin of these elements, or how have they evolved, remain a mystery. In this manuscript we report that a new family of phage satellites is able to specifically package their DNA into small-sized capsids, which are of satellite origin. Since the cf-PICI-encoded proteins responsible of these processes are similar to those encoded by their helper phages, it is tempting to speculate that they have a phage origin and have evolved not just to be able to produce the small-sized capsids, but also to avoid interaction with the phage-encoded machinery.

In most of the previously analysed satellites, the production of the small-sized capsids and the preferential packaging of the satellites are two independent processes. Thus, some SaPIs that use the *cos* system for packaging, as well as the P4 satellite, are able to produce small capsids, but they do not package their DNA preferentially, since they carry the helper *cos* sequence in their genomes and use the phage terminase complex to initiate packaging (Carpena et al., 2016; Quiles-Puchalt et al., 2014; Christie and Dokland, 2012; Lindqvist et al., 1993). Another example corresponds to the SaPIs that use the headful mechanism for packaging (*pac* SaPIs). Here, both the phage- and the SaPI-encoded small terminase can interact with both the small and the large-sized capsids. Therefore, the production of the small sized capsids limits phage reproduction not because the phage TerS cannot use these small capsids, but because the phage genome cannot be completely packaged in these capsids (Ubeda et al., 2007). An additional example refers to the other family of PICIs present in *E. coli*. Although they are 1/3 of the size of their helper phages, these elements are packaged in large capsids (probably carrying 3 concatemeric molecules per capsid), but their preferential packaging is driven by the expression of the PICI-encoded Rpp protein, which interacts with the phage TerS to modify its specificity, promoting PICI packaging (Fillol-Salom et al., 2019). Lastly, the recently described PLE satellites also produce small capsids, although the mechanism of small capsid production and how they preferentially package their genomes into these small sized capsids is still unknown (Netter et al., 2021).

Contrary to the previously described satellites, in the cf-PICI family of satellites both the production of the small-capsids and the exclusive packaging of the elements are linked. In the system characterised here, phage and satellite packaging are clearly independent, and they have evolved to be also mutually exclusive. Why do different satellites use different strategies? Most of the previously characterised satellites hijack the machinery of helper phages that use the classical *cos* or *pac* systems for packaging. In these systems, the terminase complex, composed by the phage or PICI-encoded TerS, and the phage-encoded TerL proteins, interact with their cognate *pac* or *cos* site to initiate packaging(Fillol-Salom et al., 2018)(Fillol-Salom et al., 2020). Interestingly, the new family of satellites described here hijacks a different family of phages, present both in Proteobacteria and Firmicutes, which to initiate packaging require not just the TerS-TerL complex but also the activity of an additional player, the HNH protein (Kala et al., 2014; Quiles-Puchalt et al., 2014). One can hypothesise that these specific requirements for packaging have selected for the existence of this new lineage of satellites, which have evolved to promote the formation of the small-sized capsids and their preferential packaging against these helper phages by incorporating and differentiating the phage-packaging module into the satellite genome.

Another interesting characteristic of the EcCIEDL933 element is that it does not interfere, at least in the conditions tested in these studies, with the reproduction of their helper phages. Since EcCIEDL933 is the only element of this new lineage characterised so far, we do not know if this behaviour is exclusive of this satellite or is a characteristic of this new family. Since we have previously demonstrated that by interfering with phage reproduction PICIs drive phage evolution and ecology (Frígols et al., 2015), the lack of interference observed here opens new avenues by which phages and satellites may interact. These are currently under study, although one might speculate that encoding a packaging system and a capsid may allow cf-PICIs to use the cell resources more efficiently than the other PICIs, and thus alleviate the cost of their production on the replicating helper phage. Indeed, other satellites require the production of proteins to change capsid size and some of them cannot preclude the (non-productive) partial packaging of phages in the small capsids. The cf-PICIs may ensure an optimal use of resources to produce their PICI particles and have a negligible impact on the production of viral particles.

Importantly, this new strategy of parasitism is present in satellites from Proteobacteria and Firmicutes, confirming its importance in nature. Their phylogeny and the wGRR analyses show that they are clearly distinct from phages, even for the proteins that have homologs in both types of elements. The phylogenetic analysis further shows that there are at least three groups of cf-PICIs that emerged independently raising interesting questions in relation to their origin. We propose here that the capsid and packaging genetic module were incorporated together into ancestral PICIs present in different bacterial species and obtained from different phages. Given the large evolutionary distances between cf-PICI and phages even in the highly conserved large subunit of the terminase, the acquisition of the phage capsid and packaging modules must have occurred a long time ago. The multiple independent acquisition of the modules from different phages explains the phylogenetic patterns and our observation of different genetic organisation in the different groups. These modules then evolved to adapt to the cf-PICI biology, which may have resulted in the elimination of accessory genes to fit the small cf-PICI genomes leading to a minimal capsid module that has remarkably similar types of components in the three groups. These components may correspond to a minimal set of genes necessary to build a capsid.

Once again, PICIs have provided unexpected insights in the biology of the satellite viruses. Because of the simplicity of the experimental models, evolutionary studies with phages and their satellites can give insight into the general principles of the biology of the eukaryotic viruses and their satellites. We have applied this concept here and have demonstrated the existence of a novel way by which satellites and helper phages interact. We anticipate that a better characterisation of this interaction will contribute to our understanding on how viruses and satellites evolve, and how this evolution and coexistence impacts on the biology of both the prokaryotic and eukaryotic cells infected by these elements.

## Methods

### Bacterial strains and growth conditions

Bacterial strains, plasmids and oligonucleotides used in this study are listed in Table S5, Table S6 and Table S7, respectively. Strains were grown at 37°C or 30°C on Luria-Bertani (LB) agar or in LB broth with shaking (120 r.p.m.) supplemented with Ampicillin (100 µg ml^-1^), Kanamycin (30 µg ml^-1^) or Chloramphenicol (20 µg ml^-1^; all Sigma-Aldrich) when required.

### Plasmid construction

The plasmids used in this study (Table S6) were constructed by cloning PCR products, amplified with primers listed in Table S7, into the pBAD18 vector using digestion and ligation. Plasmids were verified by Sanger sequencing in Eurofins Genomics.

### DNA methods

The introduction of the chloramphenicol (*cat*) resistance marker into the EcCIEDL933 element was performed as described (Fillol-Salom et al., 2018) using λ Red recombinase-mediated recombination. Briefly, marker was amplified by PCR using primers listed in Table S7 from plasmid pKD3, and the PCR product was transformed into the recipient strain harbouring plasmid pRWG99, which expresses the λ Red recombinase, and the marker was inserted into the PICI genome. The insertion of the resistance marker was verified by PCR.

For PICI mutagenesis, the site-directed scarless mutagenesis was performed as described previously (Blank et al., 2011; Hoffmann et al., 2017). Briefly, the *km*R marker together with an I-*Sce*I recognition restriction site was amplified by PCR, using primers listed in Table S7. Then, PCR product was inserted into the recipient strain harbouring plasmid pRWG99, which expresses the λ Red recombinase protein. After verification of the insertion by PCR, 80mer DNA fragments derived from oligonucleotides which contains the desired mutation were electroporated into the mutant strain expressing the λ Red recombinase-mediated system, and successful recombinants were selected by expression of I-*Sce*I endonuclease. The different PICI mutants obtained were verified by PCR and Sanger sequencing in Eurofins Genomics.

For phage mutagenesis, an allelic replacement strategy was performed (Solano et al., 2009). Briefly, the allelic-exchange vector, pK03-Blue, was used to clone the desired genes using the primers listed in Table S7. Then, plasmid was inserted into the strain carrying the prophage and transformants were selected on LBA plates supplemented with chloramphenicol and incubated at 32°C for selection of the temperature-sensitive plasmid. To produce the homologous recombination, the plasmid was forced to integrate into the phage genome at the non-permissive temperature (42°C). Light blue colonies, which are indicative of plasmid integration, were grown in LB broth at 32°C and ten-fold serial dilution of the overnight cultures was plated on LBA plated containing X-gal (5-bromo-4-chloro-3-indolyl-B-D-galactopyranoside) and sucrose 5% (to force the plasmid loss) and incubated at 32°C for 24 h. White colonies, which is indicative that the plasmid is loss, were screened for chloramphenicol sensitivity. The different phage mutants obtained were verified by PCR and Sanger sequencing in Eurofins Genomics.

### Phage plaque assay

A 1:50 dilution (in fresh LB broth) of the overnight strains were grown until an OD_600_=0.34 was reached. Bacterial lawns were prepared by mixing 300 µL of cells with phage top agar (PTA) and poured onto square plates. Then, serial dilutions of phages were prepared in phage buffer (50mM Tris pH 8, 1mM MgSO_4_, 4mM CaCl_2_ and 100mM NaCl) and spotted on correspond plate, which were incubated at 37°C for 24h.

### Phage and PICI induction

For PICI and phage induction, overnight cultures of lysogenic donor strains in presence and absence of the PICI were diluted 1:50 in fresh LB broth and grown at 37°C and 120 r.p.m. until an OD_600_ of 0.2 was reached. Mitomycin C (2 mg ml^-1^) was added to induce the prophage and the induced cultures were grown at 32°C with slow shaking (80 r.p.m.). Generally, cell lysis occurred 4-5 h post-induction and the induced samples were filtered using sterile 0.2 μm filters (Minisart® single use syringe filter unit, hydrophilic and non-pryogenic, Sartonium Stedim Biotech). The number of phage or PICI particles in the resultant lysate was quantified.

### Phage titration

The number of phage particles were quantified using the tittering assay. Briefly, a 1:50 dilution (in fresh LB broth) of an overnight recipient strain was grown until an OD_600_ of 0.34 was reached. Then, recipient strain was infected using 100 µL of cells with the addition of 100 µL of phage lysate serial dilutions prepared with phage buffer (50mM Tris pH 8, 1mM MgSO_4_, 4mM CaCl_2_ and 100mM NaCl), and incubated for 5 min at room temperature. The different mixtures of culture-phage were plated out on phage base agar plates (PBA; 25 g of Nutrient Broth No. 2, Oxoid; 7g agar) supplemented with CaCl_2_ to a final concentration of 10mM. Plates were incubated at 37°C for 24h and the number of plaques formed (phage particles present in the lysate) were counted and represented as the plaque forming units (PFU/mL).

### PICI transduction

The number of PICI particles were quantified using the transduction tittering assay. Briefly, a 1:50 dilution (in fresh LB broth) of an overnight recipient strain was grown until an OD_600_ of 1.4 was reached. Then, strains were infected using 1 mL of recipient cells with the addition of 100 µL of PICI lysate serial dilutions prepared with phage buffer (50mM Tris pH 8, 1mM MgSO_4_, 4mM CaCl_2_ and 100mM NaCl), and cultures were supplemented with CaCl_2_ to a final concentration of 4.4mM before incubation for 30 min at 37°C. The different mixtures of culture-PICI were plated out on LBA plates containing the appropriate antibiotic. LBA plates were incubated at 37°C for 24h and the number of colonies formed (transduction particles present in the lysate) were counted and represented as the colony forming units (CFU/mL).

### Southern Blot

Following phage and PICI induction (mitomycin C; Sigma-Aldrich from *Streptomyces caespitosus*), one millilitre of each sample was taken at defined time points and pelleted. Samples were frozen at −20°C until all collections were accomplished. Then, samples were re-suspended in 50 μl lysis buffer (47.5 μl TES-Sucrose and 2.5 μl lysozyme [10 μg ml^-1^]; Sigma-Aldrich) and incubated at 37°C for 1h. Then, 55 μl of SDS 2% proteinase K buffer (47.25 μl H_2_O, 5.25 μl SDS 20%, 2.5 μl proteinase K [20 mg ml^-1^], Sigma-Aldrich from *Tritirachium album*) was added to the lysates and incubated at 55°C for 30 min. Then, samples were vortexed with 10 μl of 10x loading dye for 1h. Following this incubation, samples were frozen and thawed in cycles of 5 min incubation in dry ice with ethanol and in a water bath at 65°C. This was repeated three times. To separate the chromosomal DNA, samples were run on 0.7% agarose gel at 30V, overnight. Then, the DNA was transferred to Nylon membranes (Hybond-N 0.45 mm pore size filters; Amersham Life Science) using standard methods. DNA was detected using a DIG-labelled probe (Digoxigenin-11-dUTP alkali-labile; Roche) and anti-DIG antibody (Anti-Digoxigenin-AP Fab fragments; Roche), before washing and visualisation. The primers used to obtain the DIG-labelled probes are listed in Table S7.

### Capsid precipitation

A large-scale induction was performed to precipitate phage and PICI particles and obtain their dsDNA. Briefly, a total of 100 ml lysate was produced by MC induction. Then, lysates were treated with RNase (1 µg ml^-1^) and DNase (10 µg ml^-1^) for 30 min at room temperature. Afterwards, 1 M of NaCl was added to the lysate and incubated 1 h on ice. After incubation, the mix was centrifuged at 11,000 x *g* for 10 min at 4°C, and phages/PICIs were mixed with 10% wt/vol polyethylene glycol (PEG) 8000 and kept overnight at 4°C. Then, phages/PICIs were precipitated at 11,000 x *g* for 10 min at 4°C and the pellet was resuspended in 1 ml of phage buffer.

### Phage and PICI DNA extraction

To extract the dsDNA, lysate from phage and PICI precipitation was treated with DNase (10 µg ml^-1^) for 30 min at room temperature. Then, lysate was combined with an equal volume of lysis mix (2% SDS and 90 μg ml^-1^ proteinaseK) and incubated at 55°C for 1 h. DNA was extracted with an equal volume of phenol:chloroform:isoamyl alcohol 25:24:1 and samples were centrifuged at 12,000 x *g* for 5 min, and the aqueous phase containing the DNA was obtained. The DNA was precipitated by 0.3M NaOAc and 2.25 volume of 100% ethanol, then pelleted at 12,000 x g for 30 min at 4°C and washed once with 1 ml of 70% ethanol. After centrifugation, the DNA pellets were air dried for 30 min and resuspended in 100 μl nuclease free water.

### Electron microscopy

To analyse the lysate by transmission electron microscopy (TEM), lysates were CsCl purified. Briefly, the precipitated phages and PICIs were loaded on the CsCl step gradients (1.35, 1.5 and 1.7 g ml^-1^ fractions) and centrifuged at 80,000 x *g* for 2h at 4°C. The phage and PICI bands were extracted from the CsCl gradients using a 23-gauge needle and syringe. Then, phages and PICIs were dialyzed overnight to remove CsCl excess using SnakeSkin^TM^ Dialysis Tubing (3.5K MWCO, 16mm dry) into 50mM of Tris pH 8 and 150mM NaCl buffer. For TEM analysis, ten microliters of the dialysed samples were incubated in a carbon-coated gold grid for 5 min. Then, samples were fixed with 1% Paraformaldehyde for 2 min, before washing three times with distilled water for 30 sec. Afterwards, the samples were stained with 2% Uranyl Formate for 30 sec, and they were allowed to dry at room temperature for 15 min. The JEOL 1200 TEM microscope was used to examine the samples. Photos were obtained at 12K of magnification.

### Identification of PICIs and KEGG analysis

The KEGG analysis performed previously(Fillol-Salom et al., 2018) suggested that the EcCIEDL933 element is the prototypical member of a new PICI family. To identify more members of this family, we searched manually on the NCBI database for elements with a size comprised between 10-15 kb, encoding a divergent pair or transcriptional regulators next to an *int* gene (Martínez-Rubio et al., 2017), and a packaging module similar to that present in EcCIEDL933. Additionally, these elements should have unique attachment sites that are never occupied by prophages, and they should lack phage lytic and tail genes. This search was followed by the analysis of orthologues performed for the EcCIEDL933 element (Table S1), which corroborates that the newly identified elements correspond to PICIs. The ortholog analysis was performed using the Kyoto Encyclopedia of Genes and Genomes (KEGG) databse (http://www.genome.jp/kegg; release July 12, 2018).

### Phylogenetic analyses

We fetched the protein sequences of TerL and the Capsid from all cf-PICI in Fig. S2 and S6 (Table S2 and S4). For each protein we searched for homologs among a database of all (3725) complete phages present in NCBI RefSeq (last accessed March 2021), using Diamond v2.0.6.144 (Buchfink et al., 2015) with parameters *--query-cover 50 --ultra-sensitive -- forwardonly*. We picked hits in phages that match either a capsid or a TerL of any cf-PICI with an e-value of at most 1e^-10^, and made a multiple alignment of these with the corresponding cf-PICI proteins, using mafft-linsi v7.490 (default parameters)(Katoh and Standley, 2013). The multiple alignments of TerL and Capsid were then purged from non-informative sites using clipkit v1.3.0 (Steenwyk et al., 2020) with the option *-m kpi-gappy*. We used the resulting multiple alignments to make phylogenetic reconstructions using maximum likelihood with iqtree v1.6.12 (Nguyen et al., 2015), using parameters -nt 6 -bb 1000 (to quantify the robustness of the topology) -m TEST (to find the best model). The trees were visualised with iTol v6.5.2 <u>(Letunic and Bork, 2021)</u> and the phage branches were collapsed for clarity.

### Weighted Gene Repertoire Relatedness

We searched for sequence similarity between all proteins of all phages using mmseqs2 (release 13-45111 (Steinegger and Söding, 2017)) with the sensitivity parameter set at 7.5. The results were converted to the blast format and we kept for analysis the hits respecting the following thresholds: e-value lower than 0.0001, at least 10% identity, and a coverage of at least 50% of the proteins. The hits were used to retrieve the bi-directional best hits between pairs of phages, which were used to compute a score of gene repertoire relatedness weighted by sequence identity:

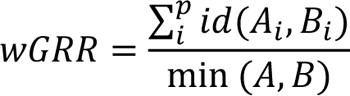

where A_i_ and B_i_ is the pair *i* of homologous proteins present in *A* and *B*, id(*Ai*,*Bi*) is the sequence identity of their alignment, and min(*A*,*B*) is the number of proteins of the genome encoding the fewest proteins (*A* or *B*). wGRR is the fraction of bi-directional best hits between two elements weighted by the sequence identity of the homologs. It varies between zero (no bi-directional best hits) and one (all genes of the smallest genome have an identical homolog in the largest genome). wGRR integrates information on the frequency of homologs and sequence identity. For example, when the smallest genome has 10 proteins, a wGRR of 0.2 can result from two homologs that are strictly identical or five that have 40% identity.

## ACKNOWLEDGEMENTS

This work was supported by grants MR/M003876/1, MR/V000772/1 and MR/S00940X/1 from the Medical Research Council (UK), BB/N002873/1, BB/V002376/1 and BB/S003835/1 from the Biotechnology and Biological Sciences Research Council (BBSRC, UK), ERC-ADG-2014 Proposal n° 670932 Dut-signal (from EU), and Wellcome Trust 201531/Z/16/Z to JRP, and grants INCEPTION project (PIA/ANR-16-CONV-0005), Equipe FRM (Fondation pour la Recherche Médicale): EQU201903007835, Laboratoire d’Excellence IBEID Integrative Biology of Emerging Infectious Diseases [ANR-10-LABX-62-IBEID], and SALMOPROPHAGE ANR-16-CE16-0029 to EPCR.

## AUTHOR CONTRIBUTIONS

JRP conceived the study; NA, LM-R and AF-S conducted the experiments; JMS and EPCR performed the genomic analyses; NA, LM-R, JMS, JC, EPCR, AF-S and JRP analysed the data. EPCR, AF-S and JRP wrote the manuscript.

## DECLARATION OF INTERESTS

Authors declare no competing interests.

---

**Table S1.**
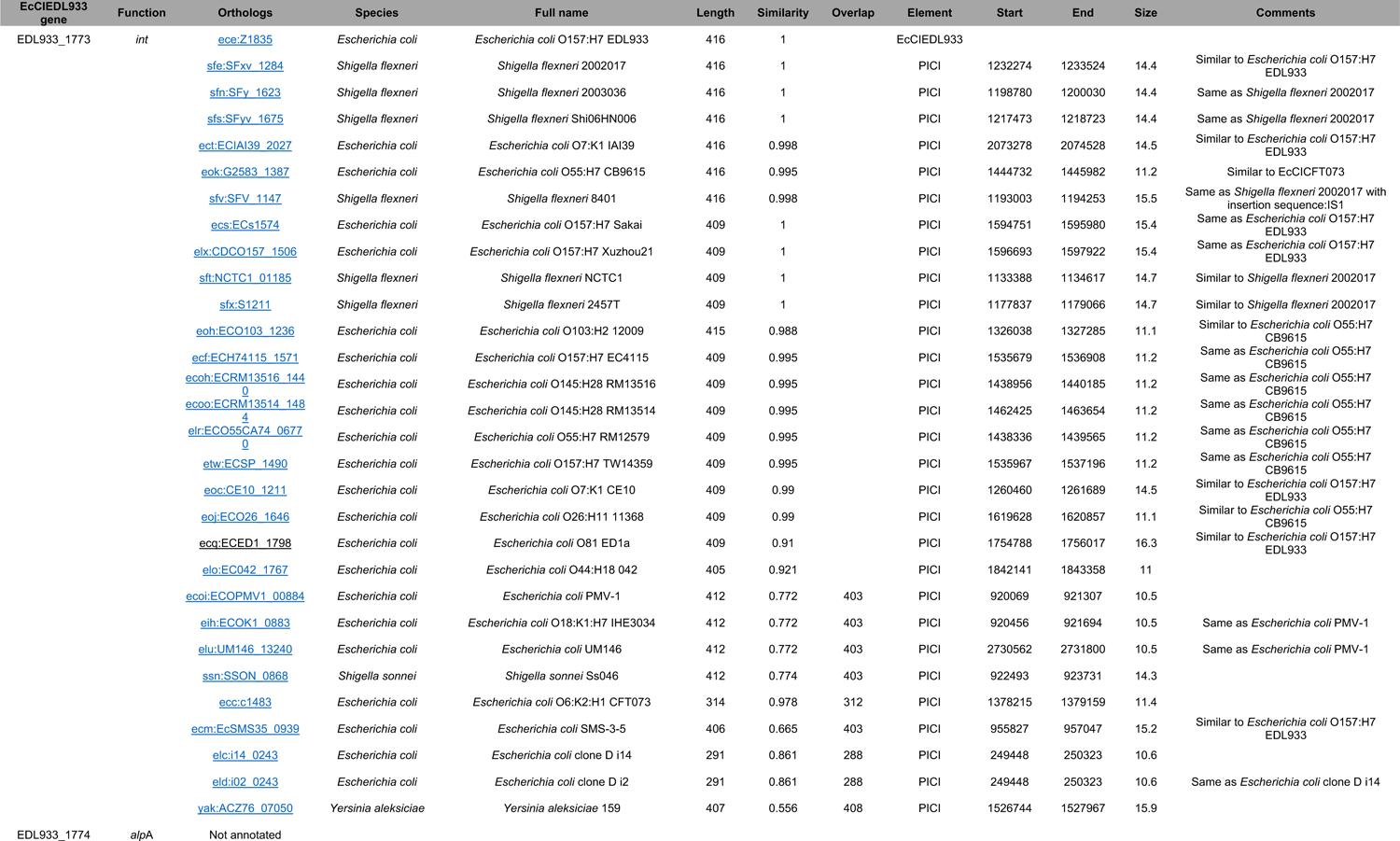
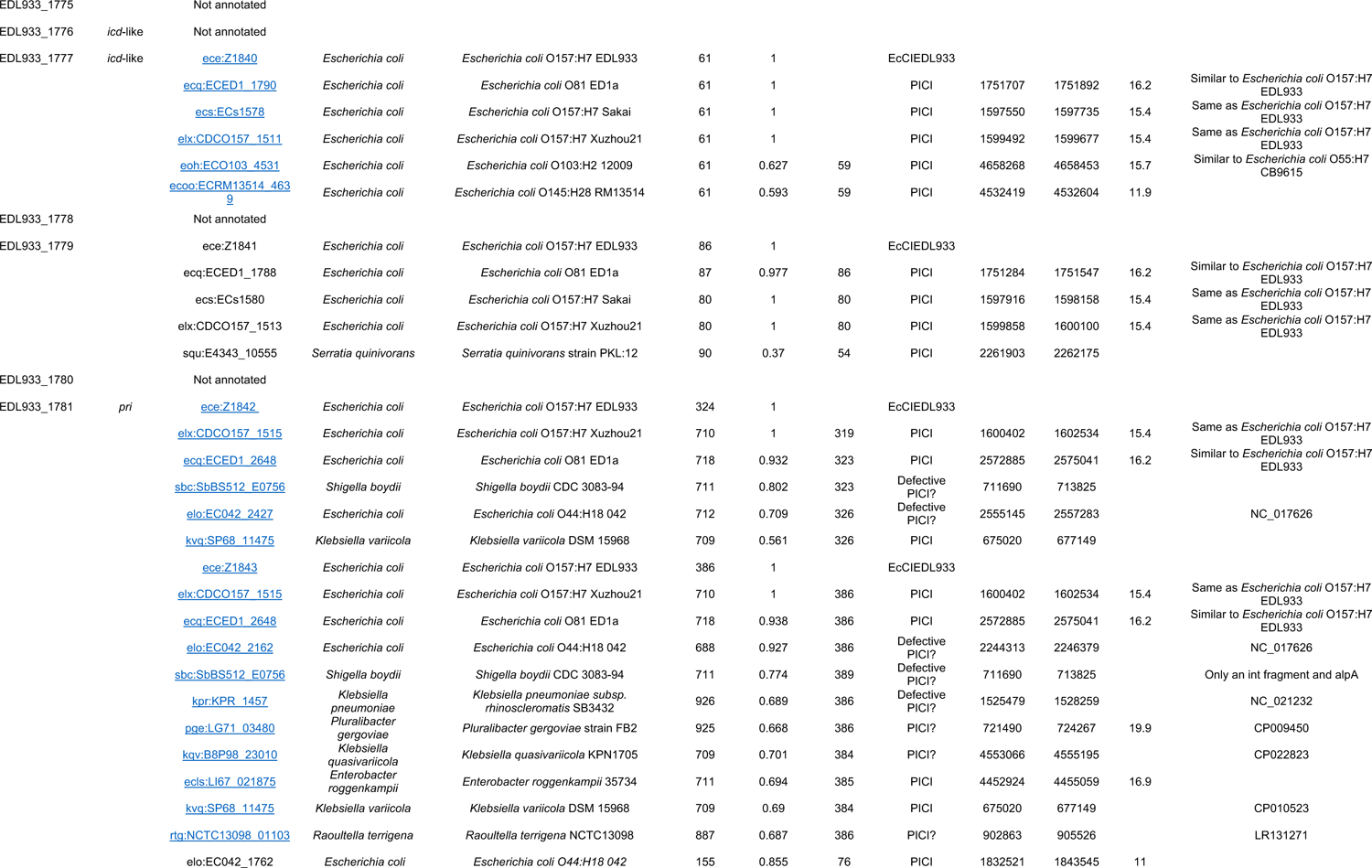
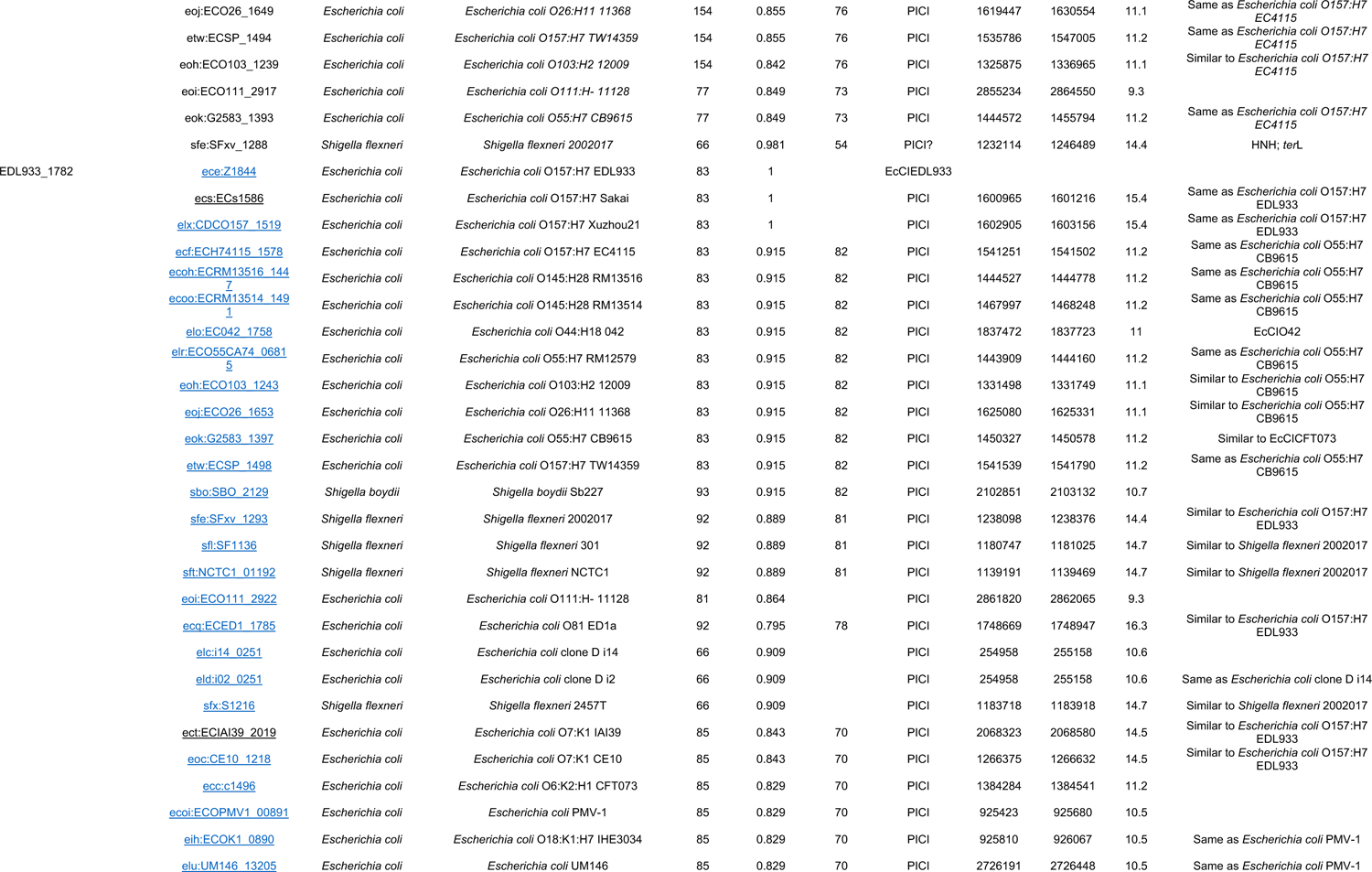
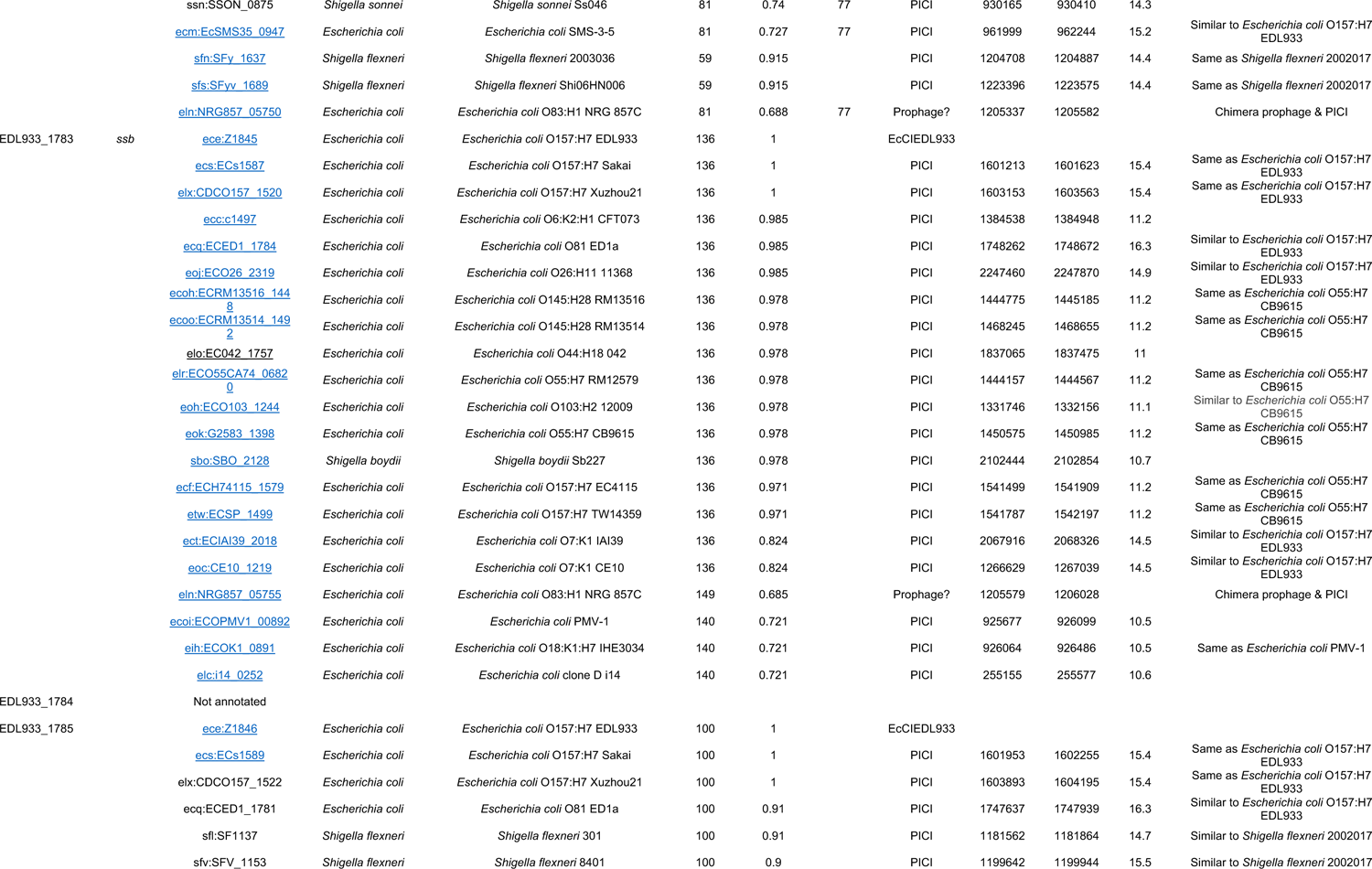
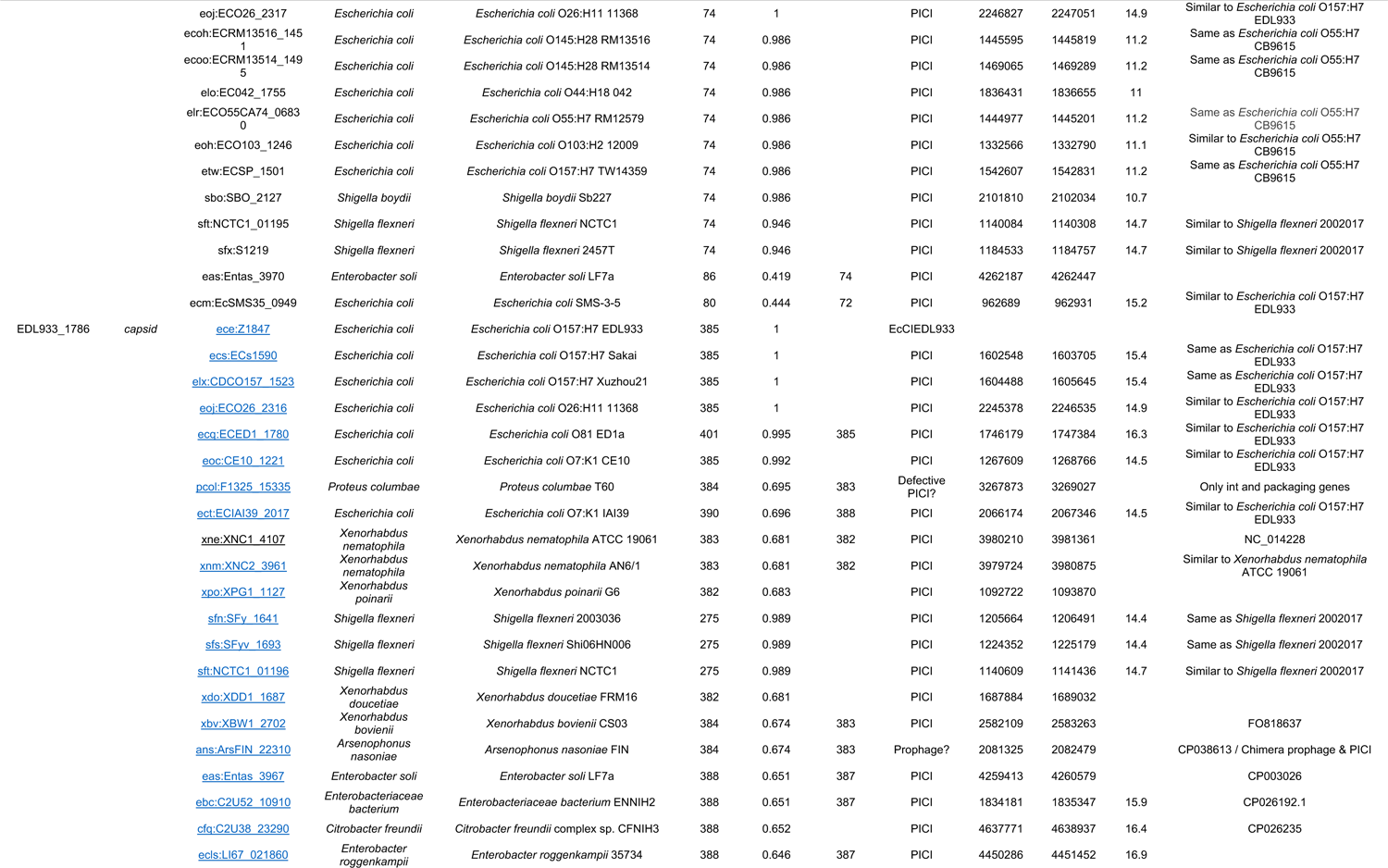
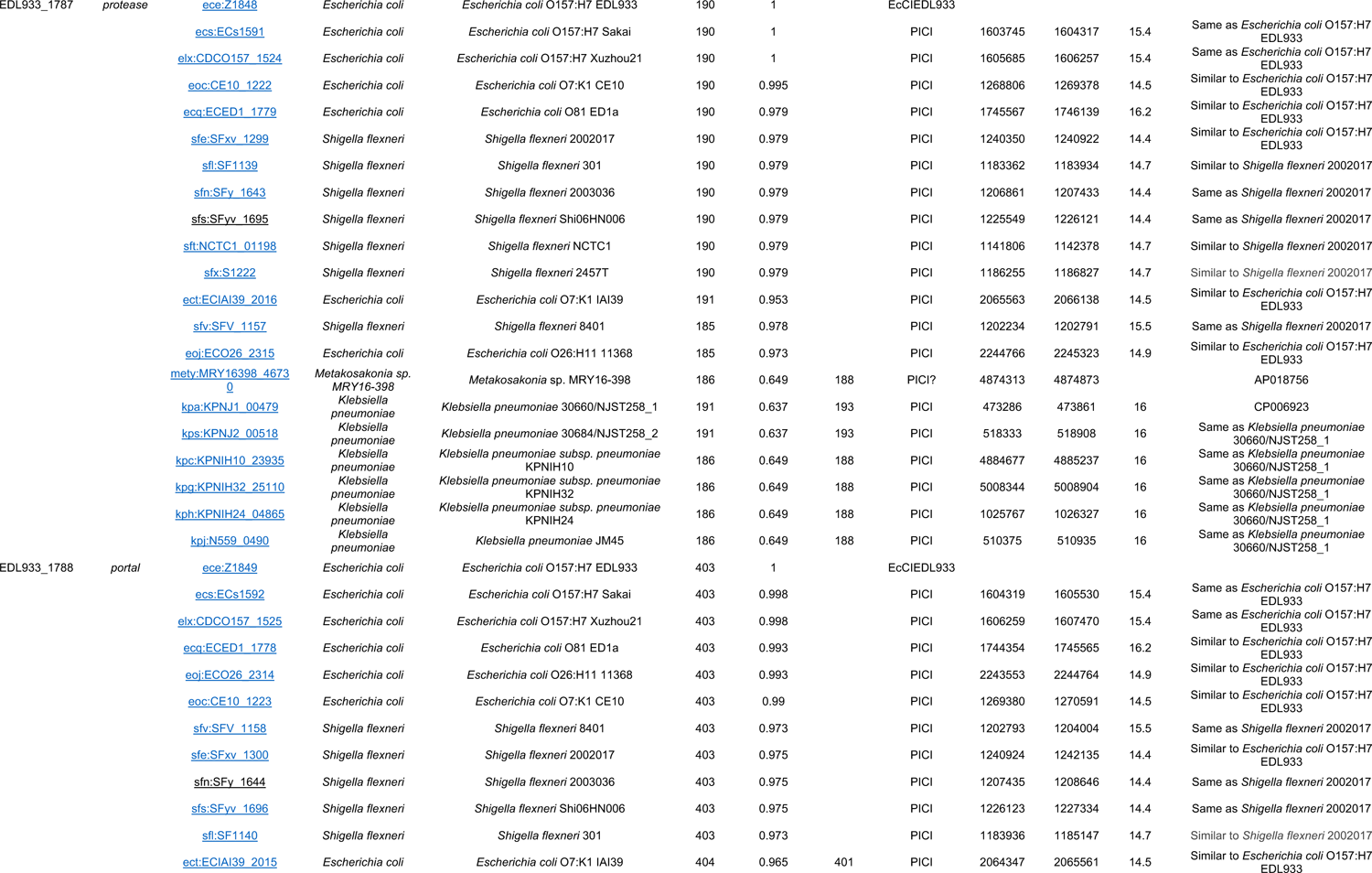
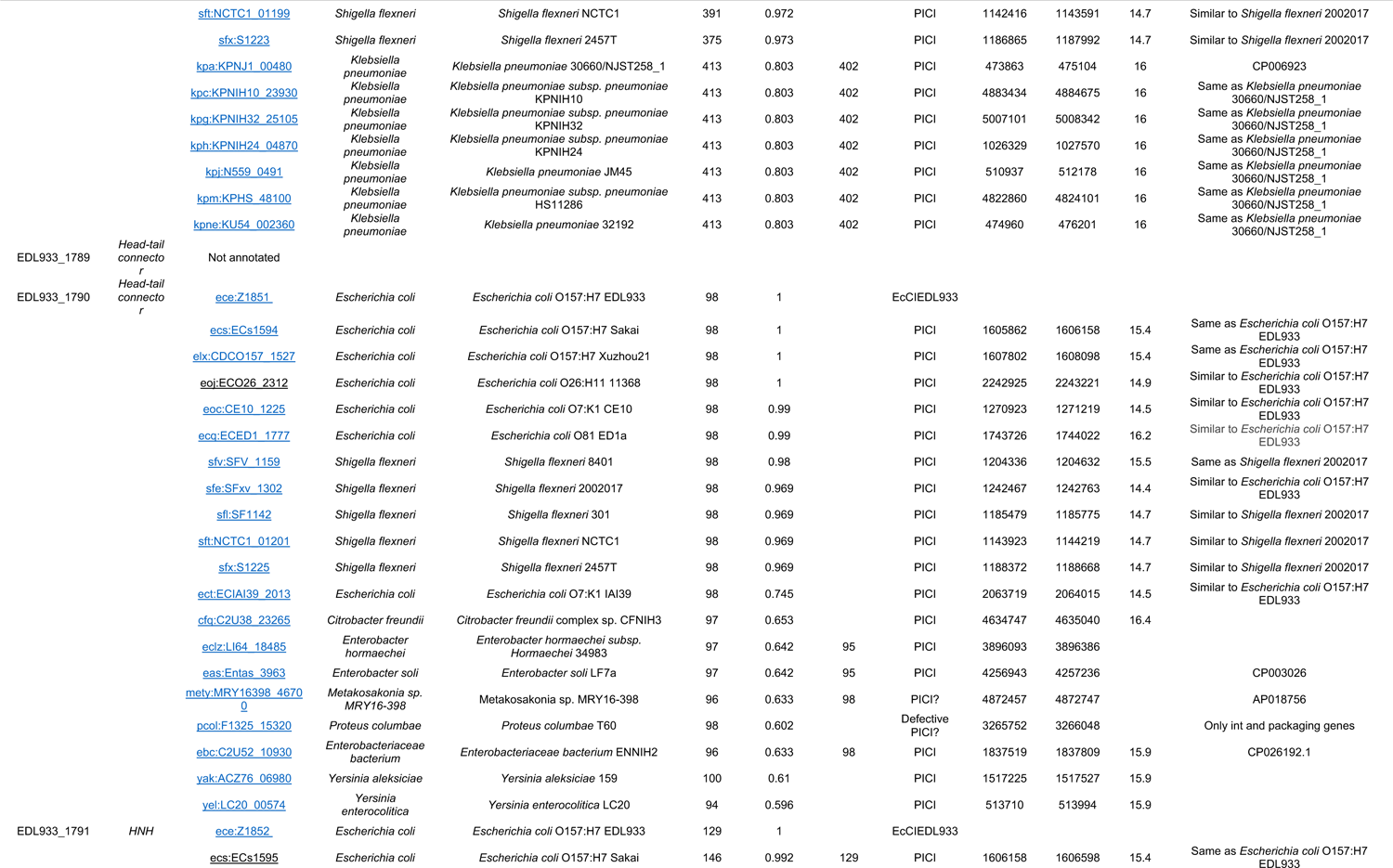
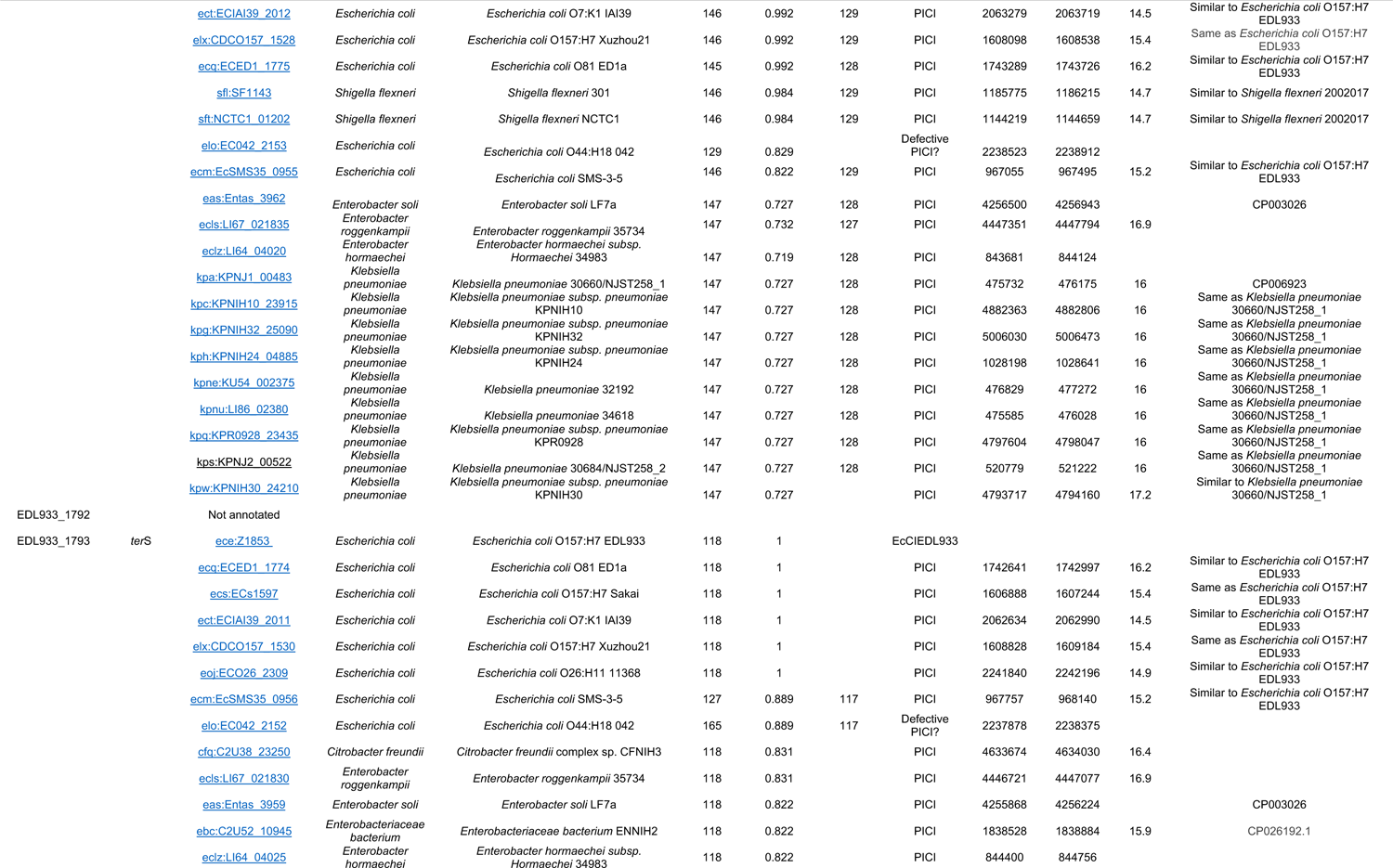
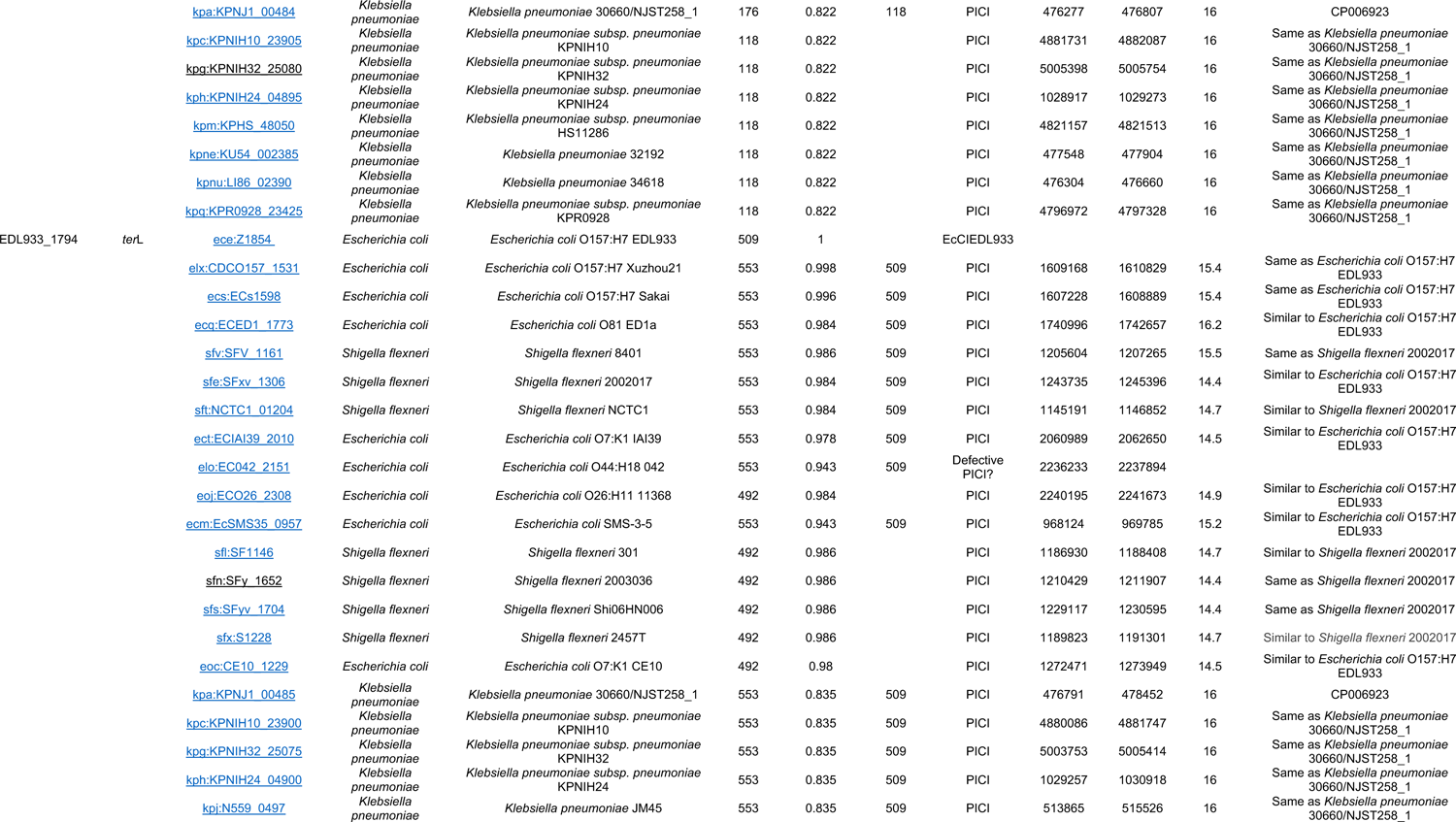
EcCIEDL933 orthologues.

**Table S2.**
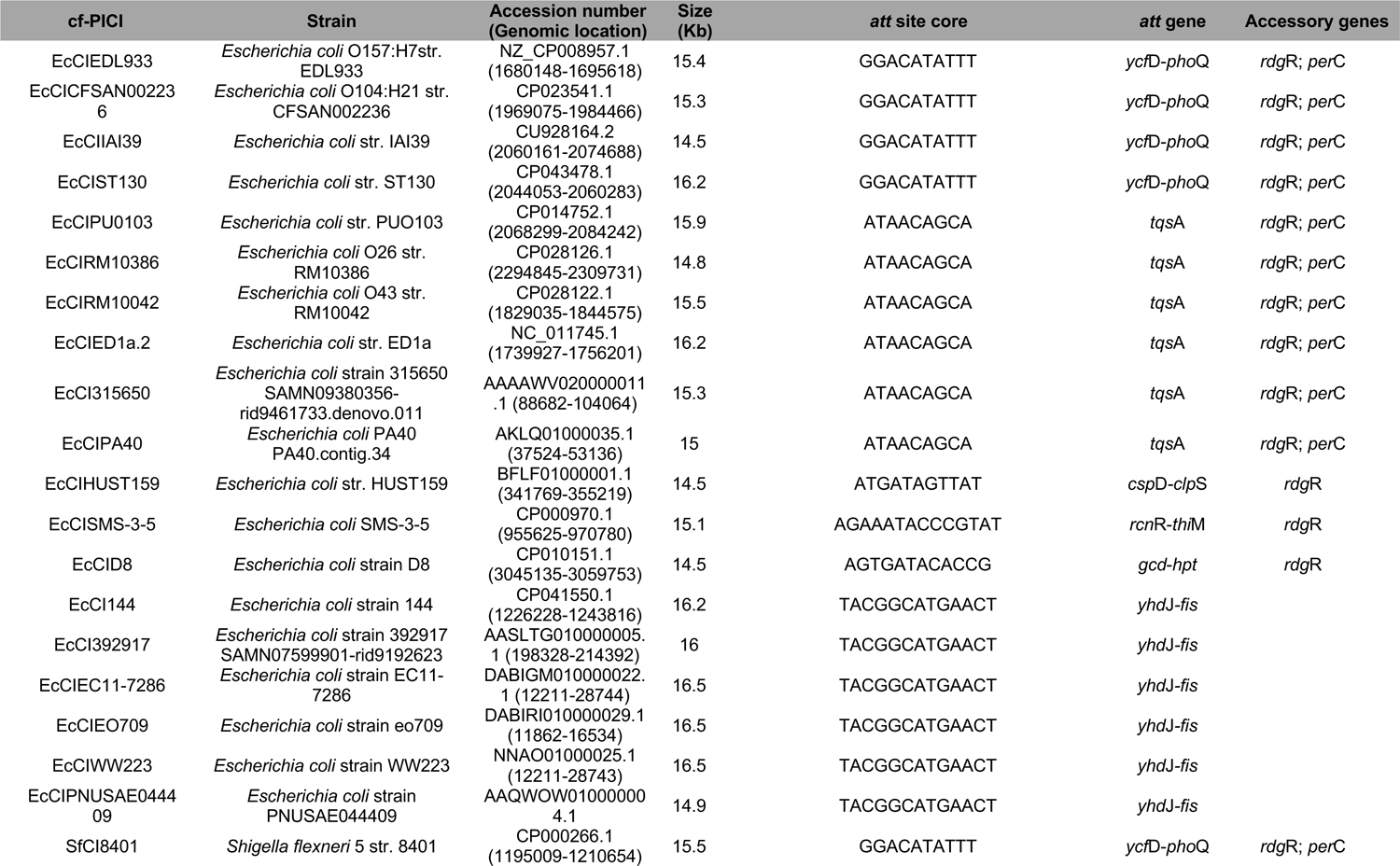
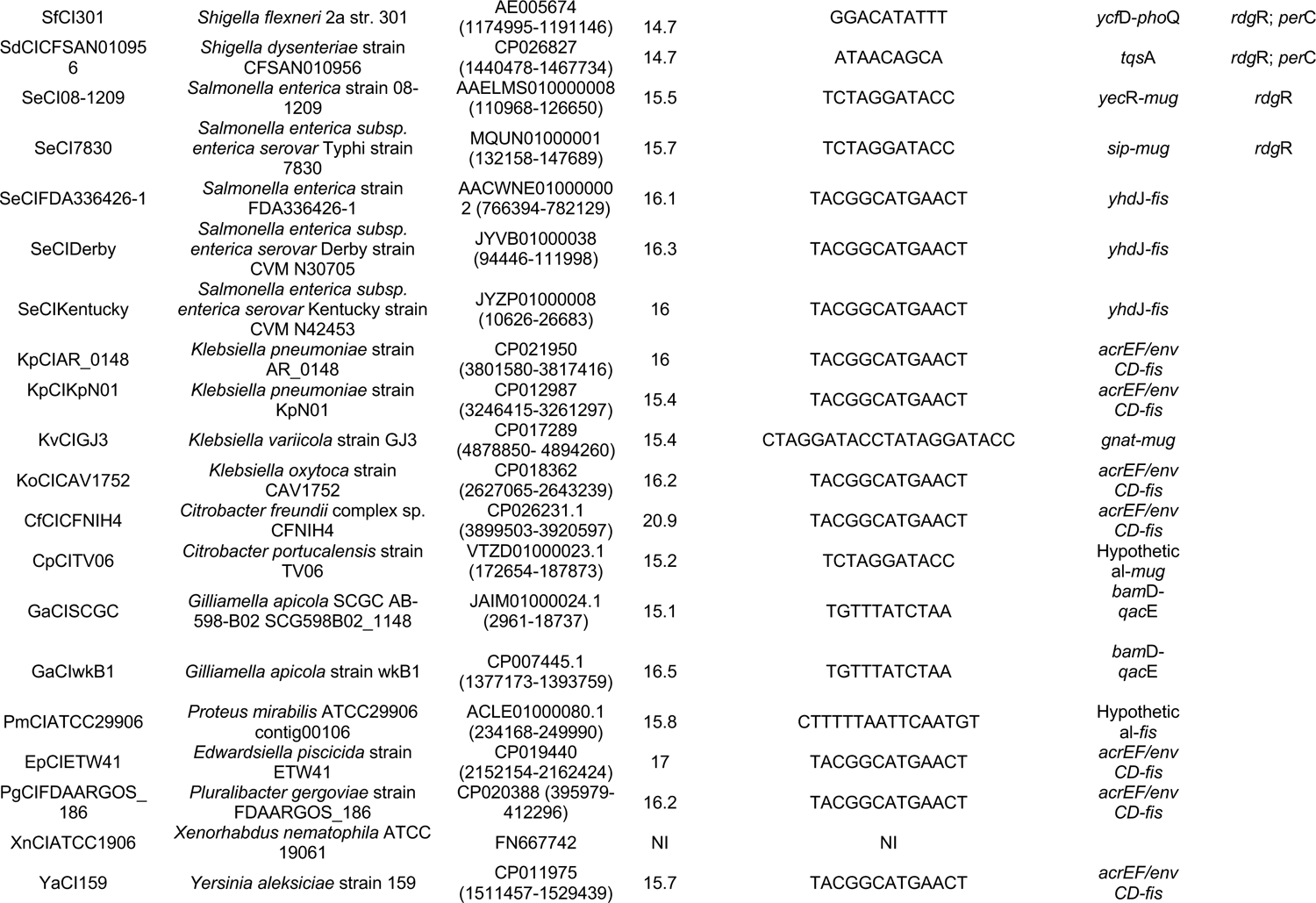
cf-PICIs in Proteobacteria: Genomes and characteristics.

**Table S3.**
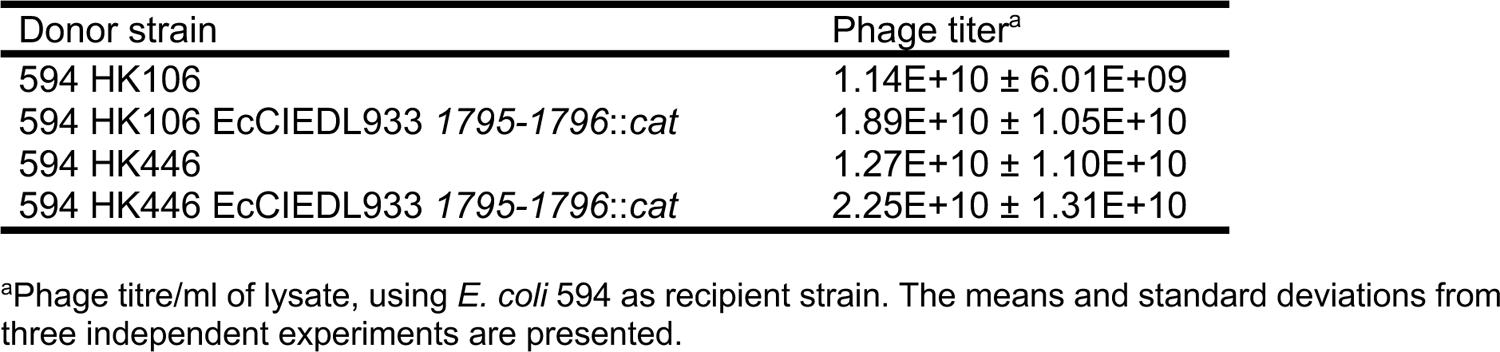
Phage titres in presence or absence of EcCIEDL933.

**Table S4.**
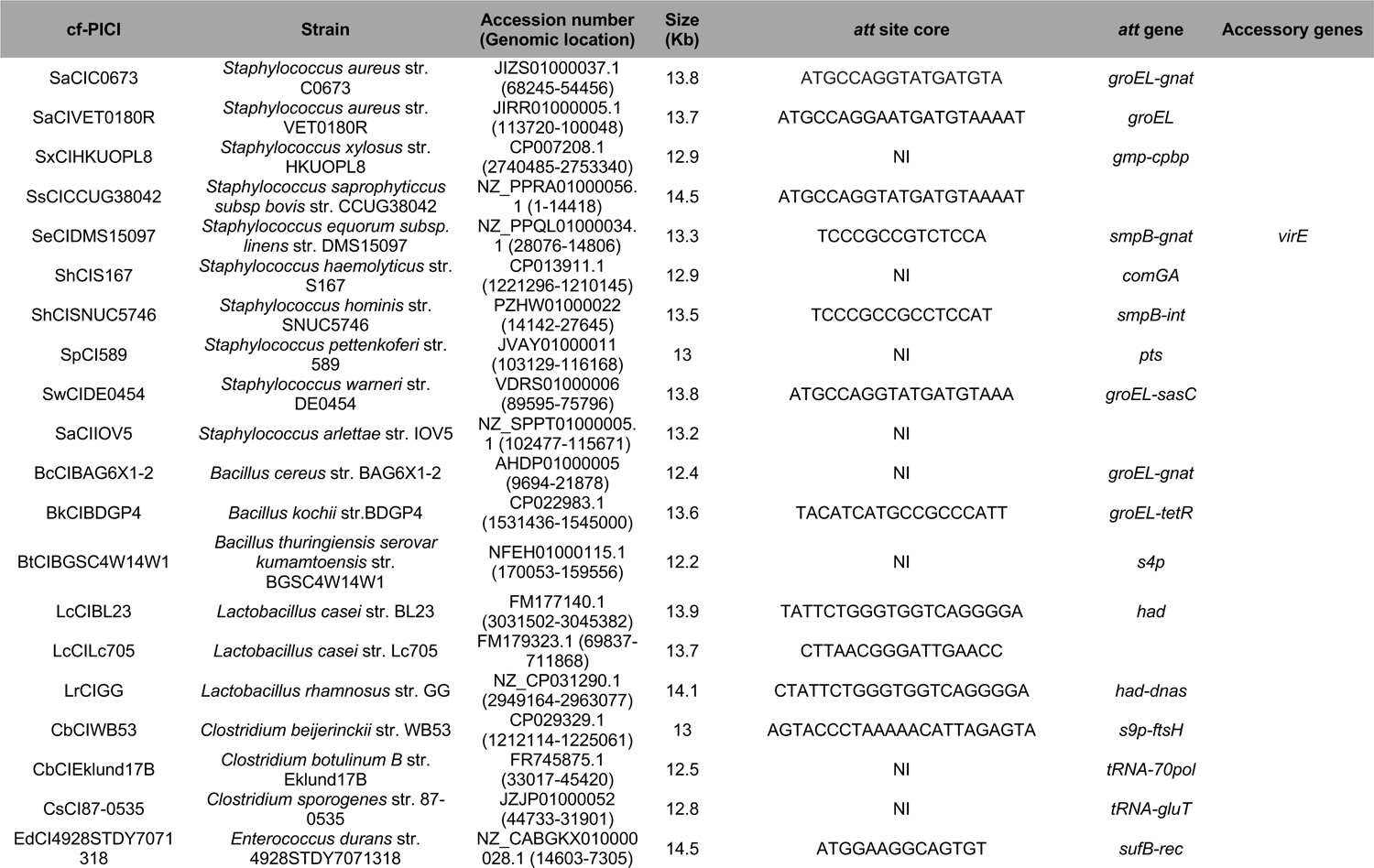
cf-PICIs in Firmicutes bacteria: Genomes and characteristics.

**Table S5.**
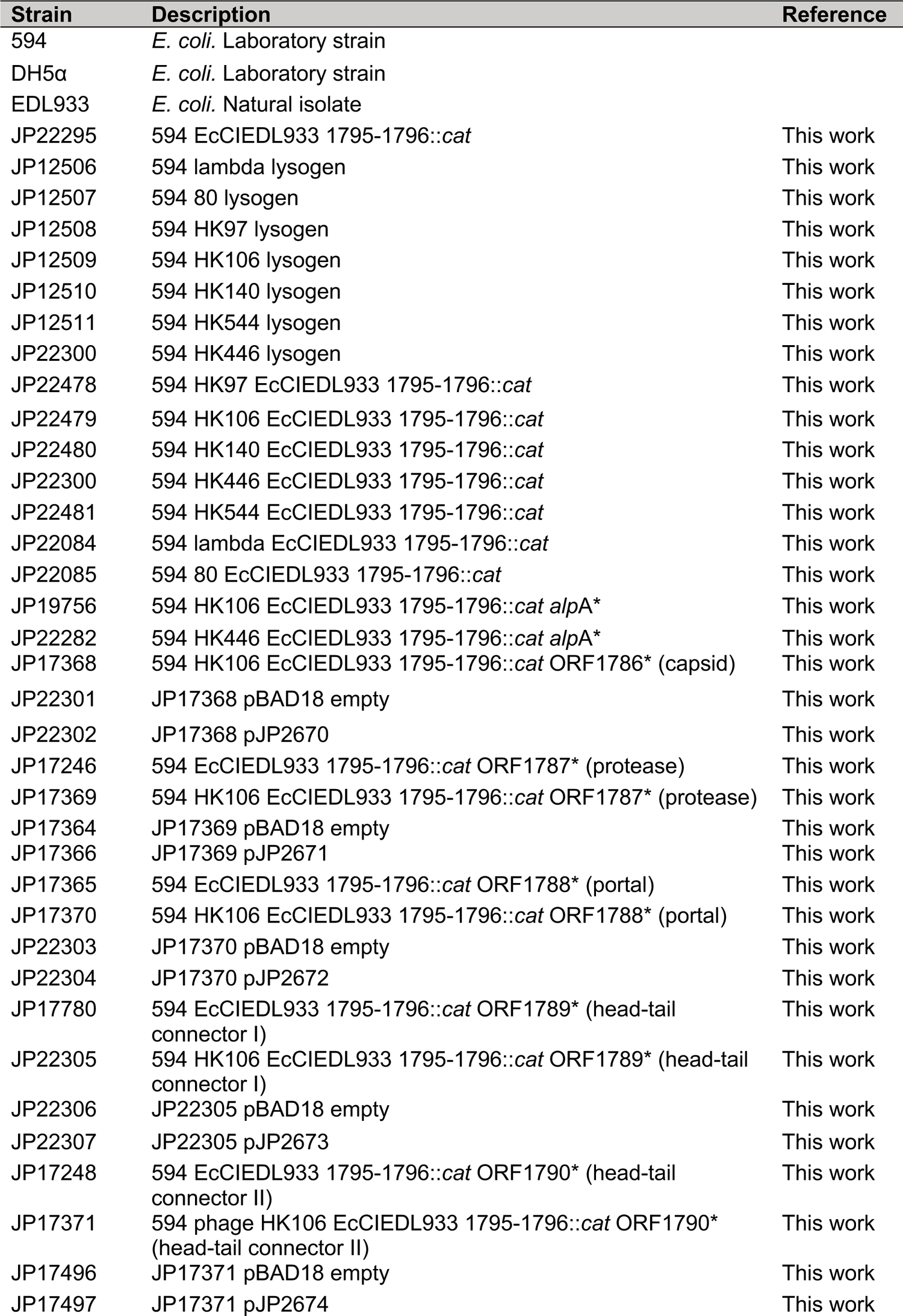
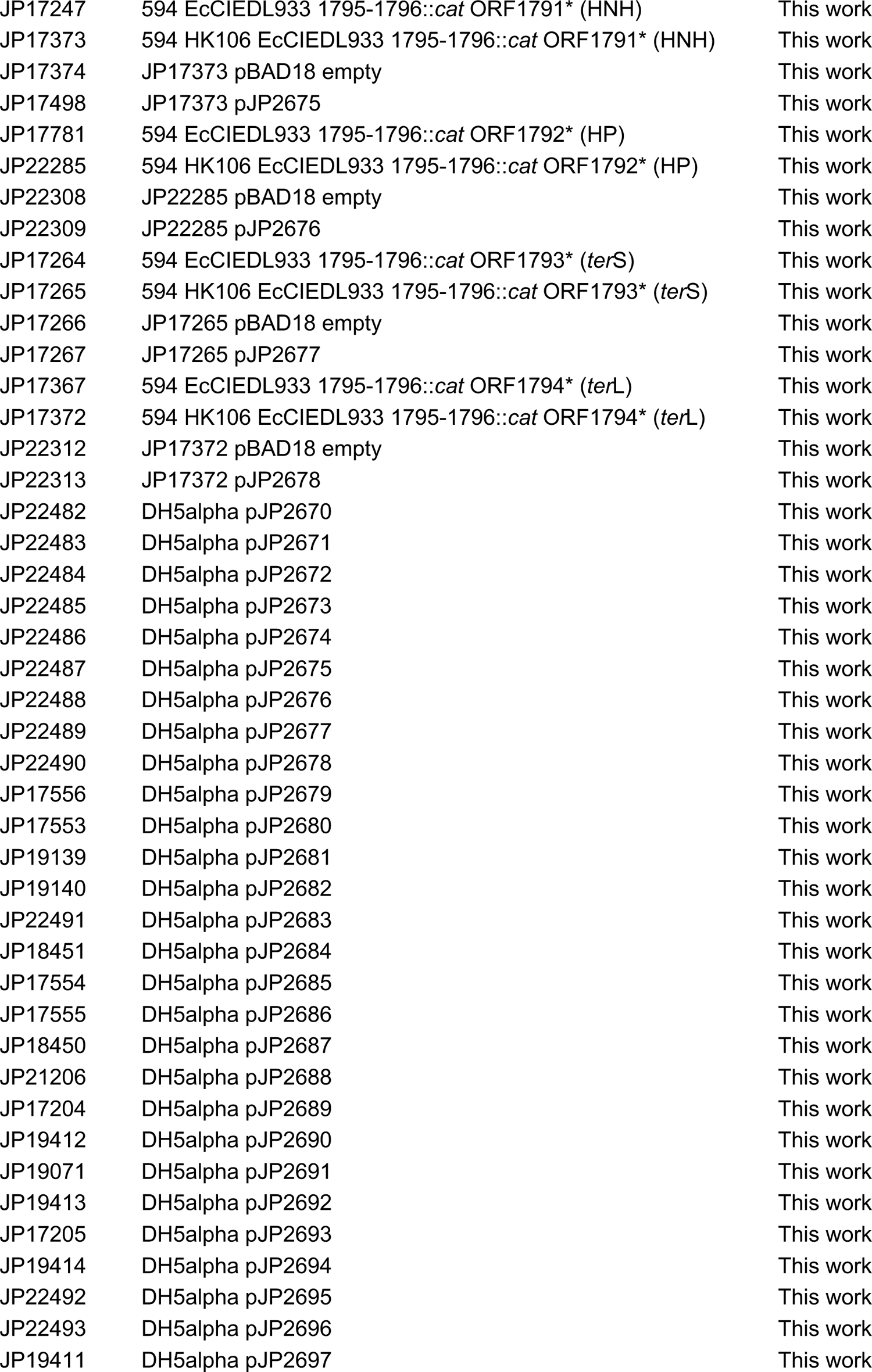
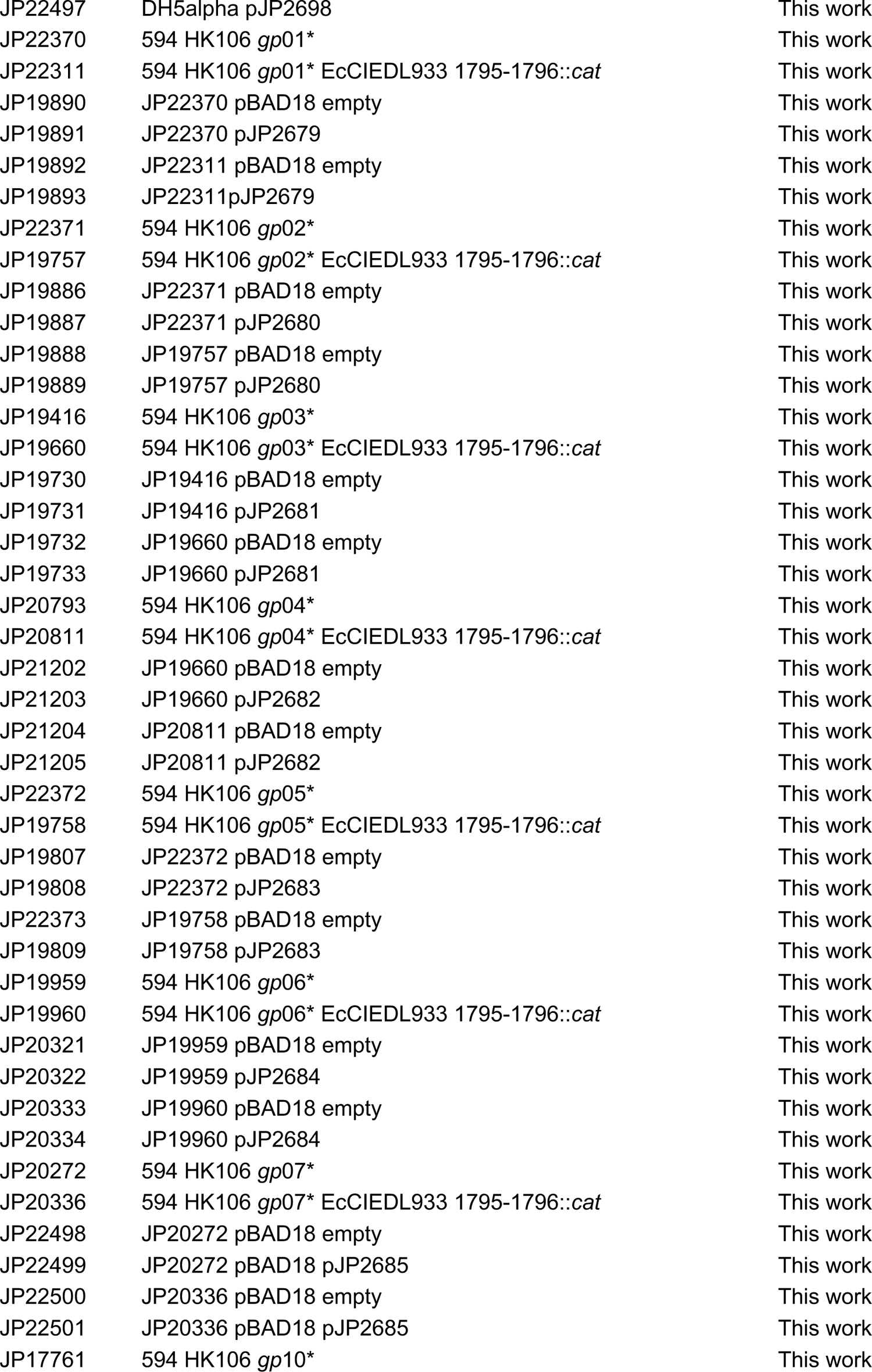
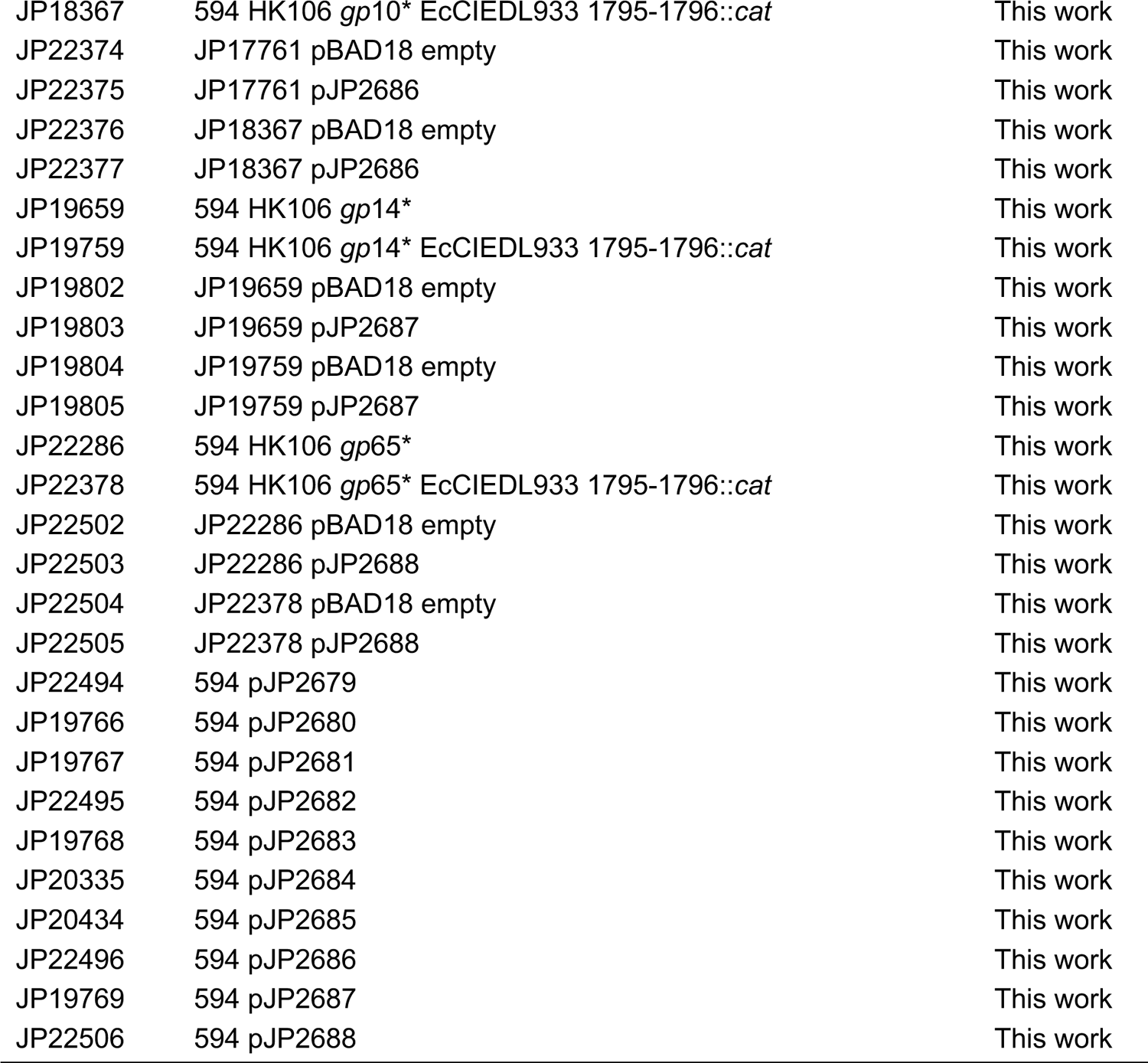
Strains used in this study.

**Table S6.**
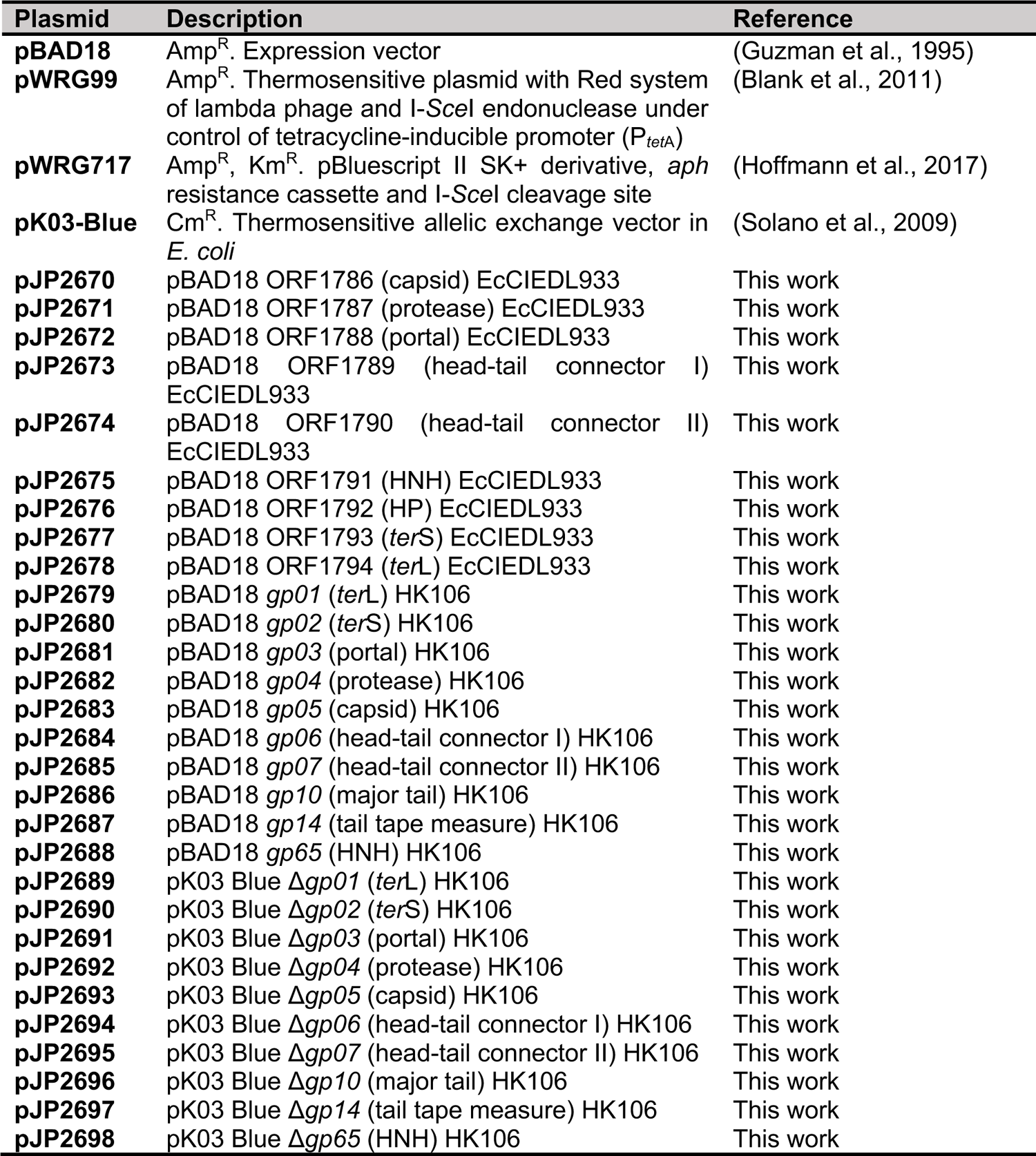
Plasmids used in this study.

**Table S7.**
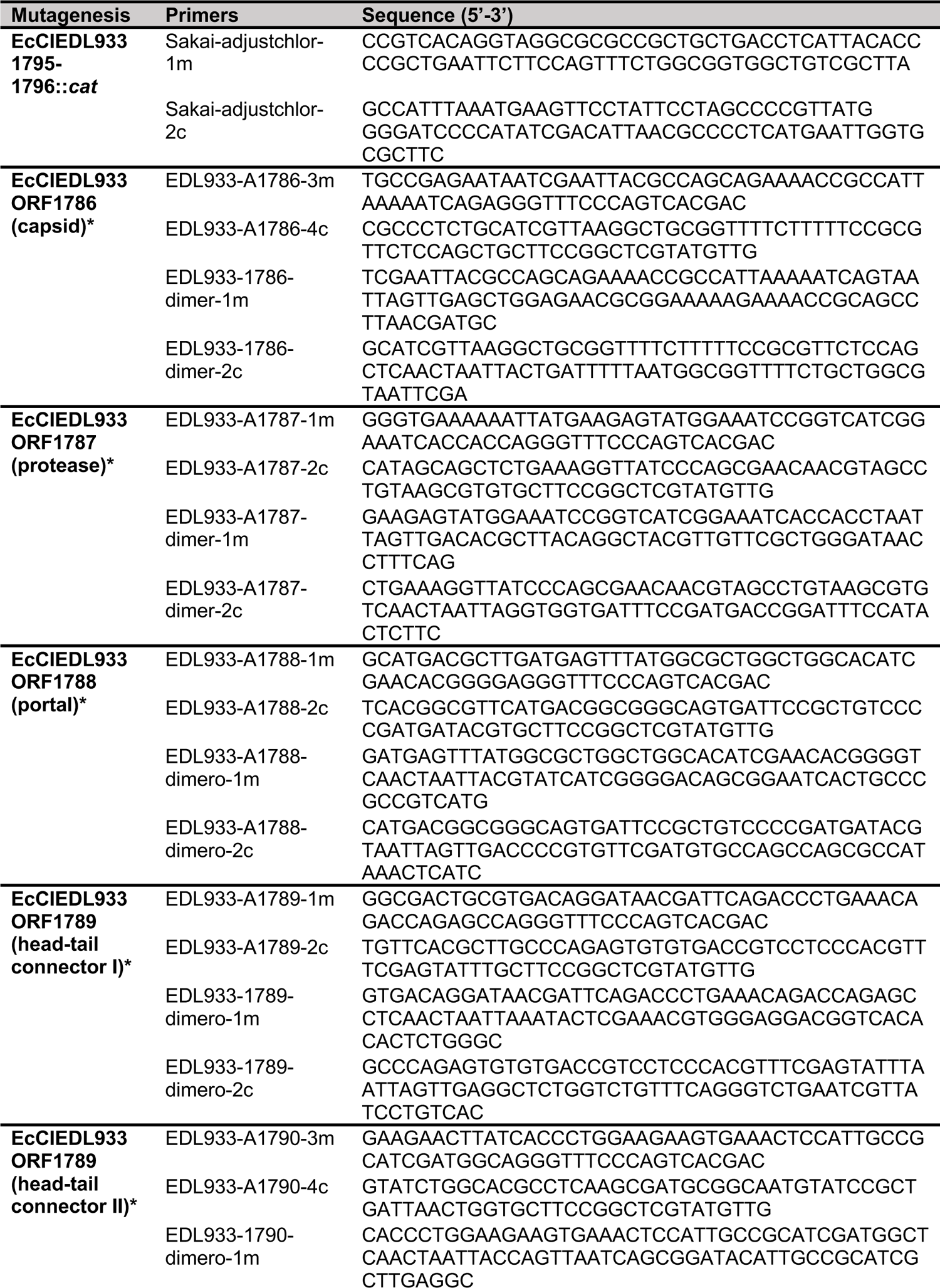
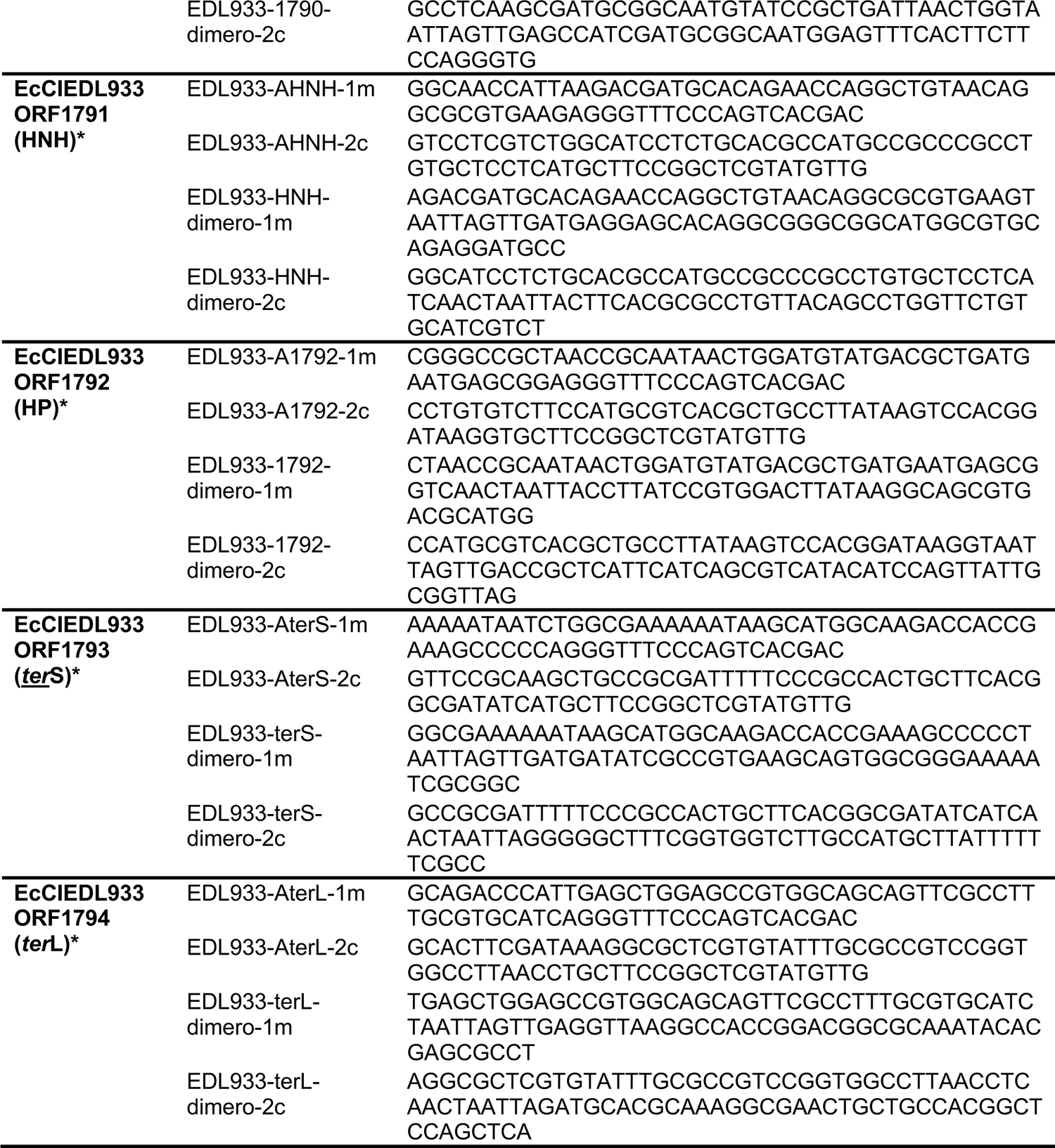
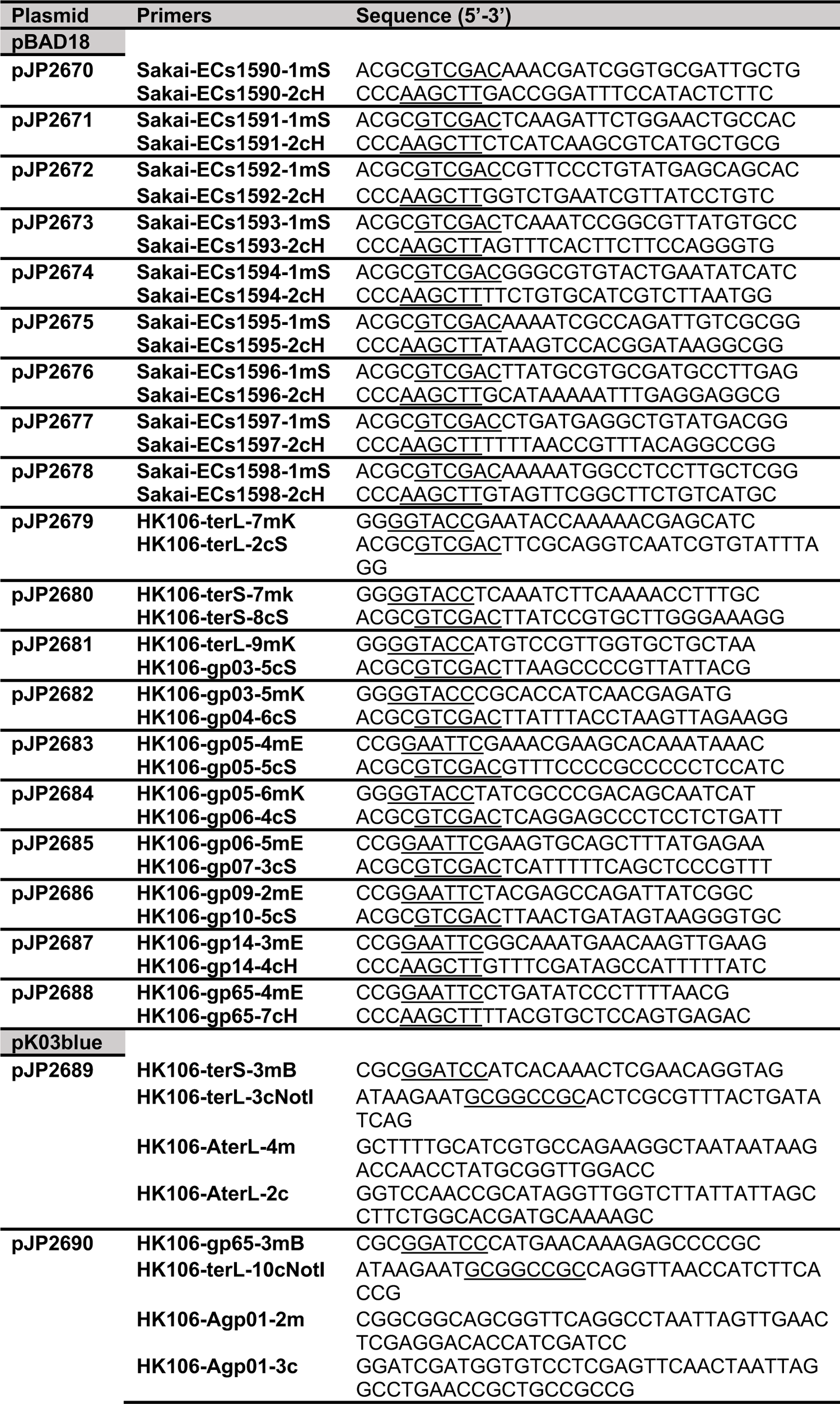
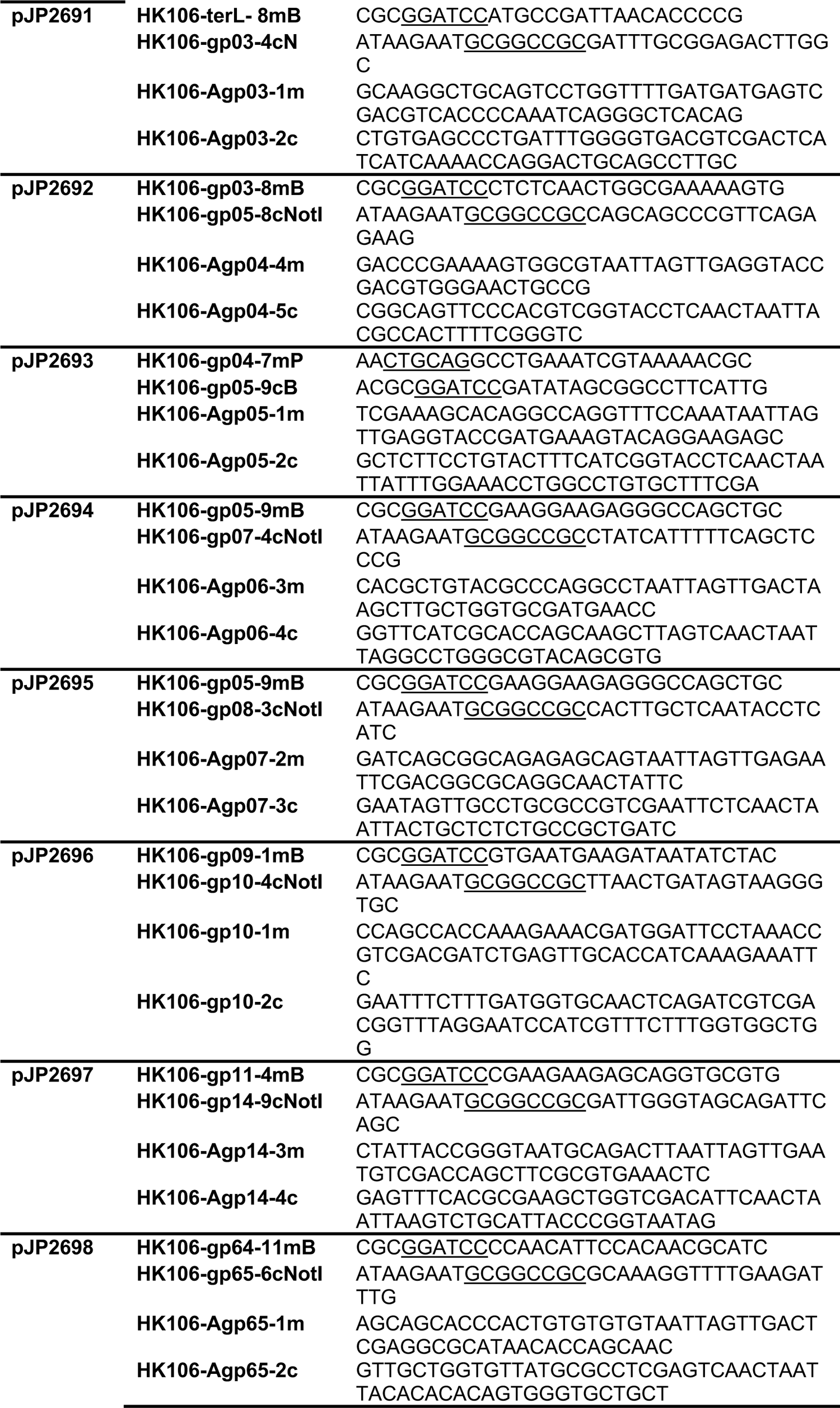
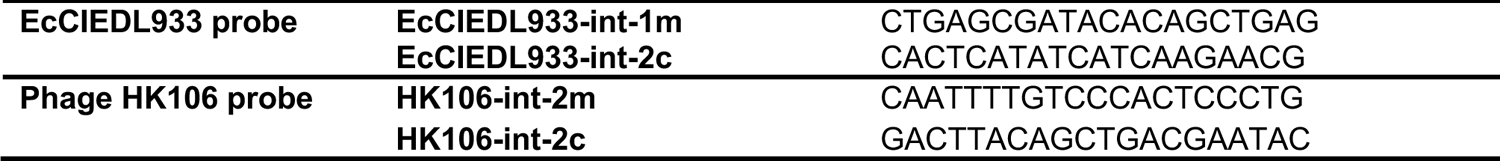
Primer list.

**Figure S1.**
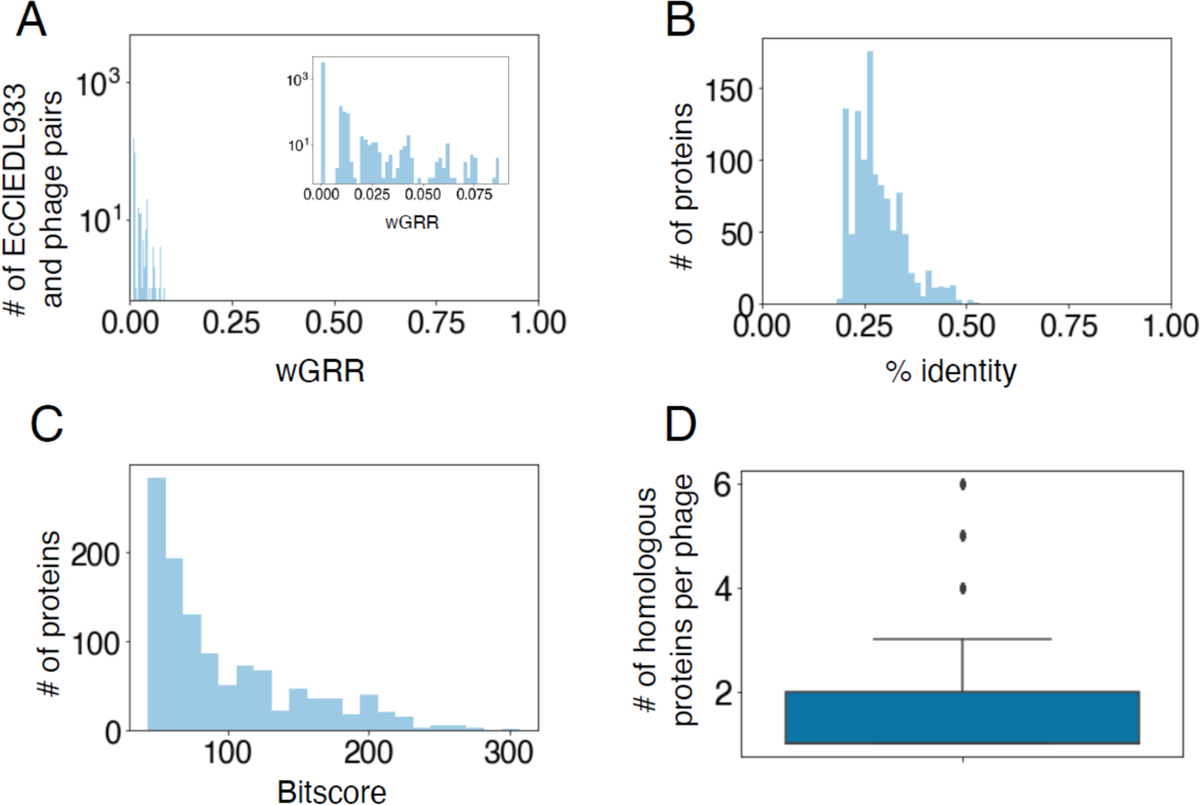
Genomic comparison between EcClEDL933 and 3725 phage genomes. (A) Distribution of wGRR values between EcClEDL933 and phage genomes. Inset shows a zoomed version between wGRR of zero and 0.1. (B) Distribution of the protein identity percentages, for all the potential homologs between EcClEDL933 and phage genomes. (C) Distribution of the bitscores for all the potential homologs between EcClEDL933 and phage genomes. (D) Boxplot with the number of homolog genes per phage, for all the phages in the database.

**Figure S2.**
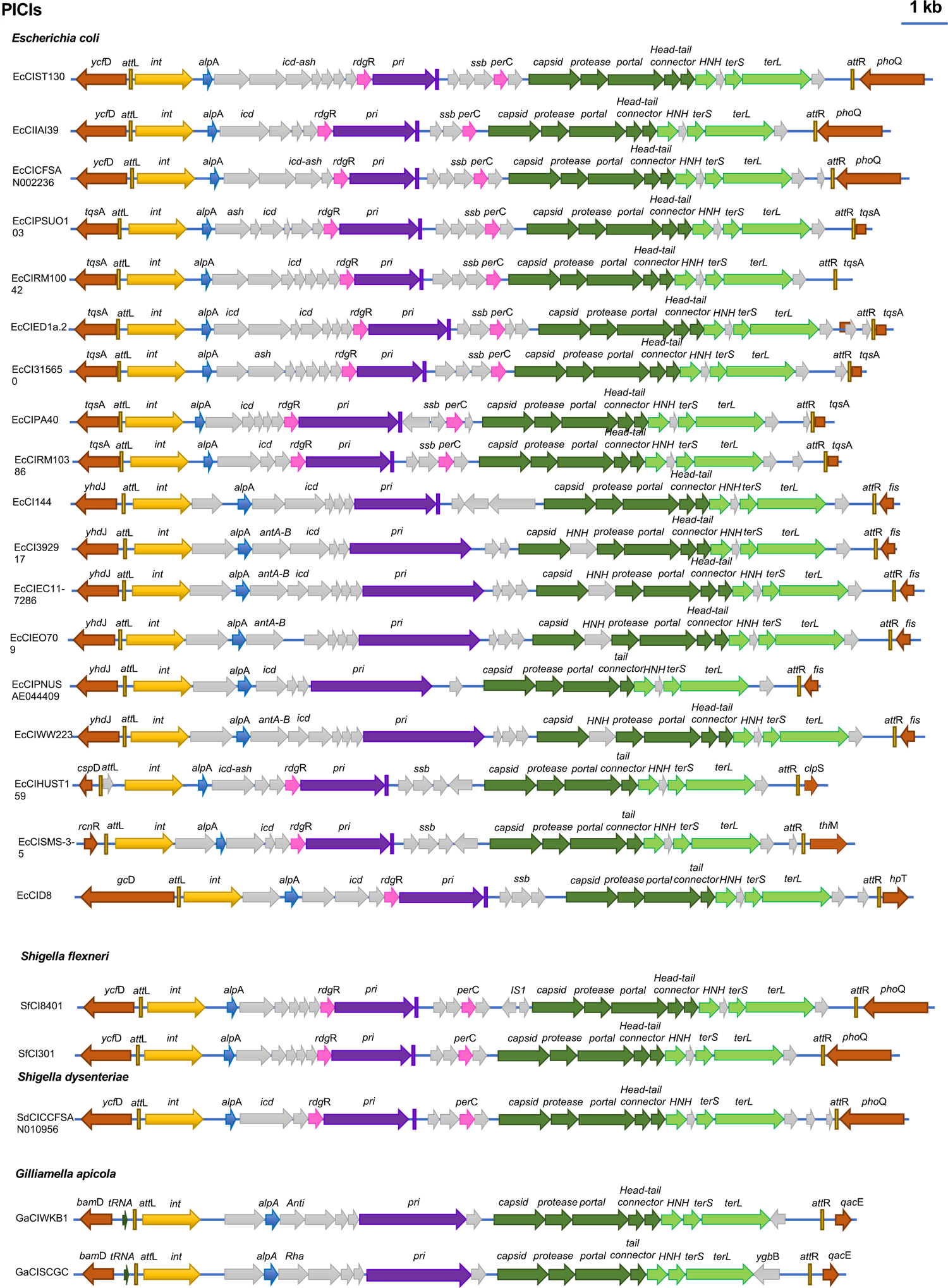
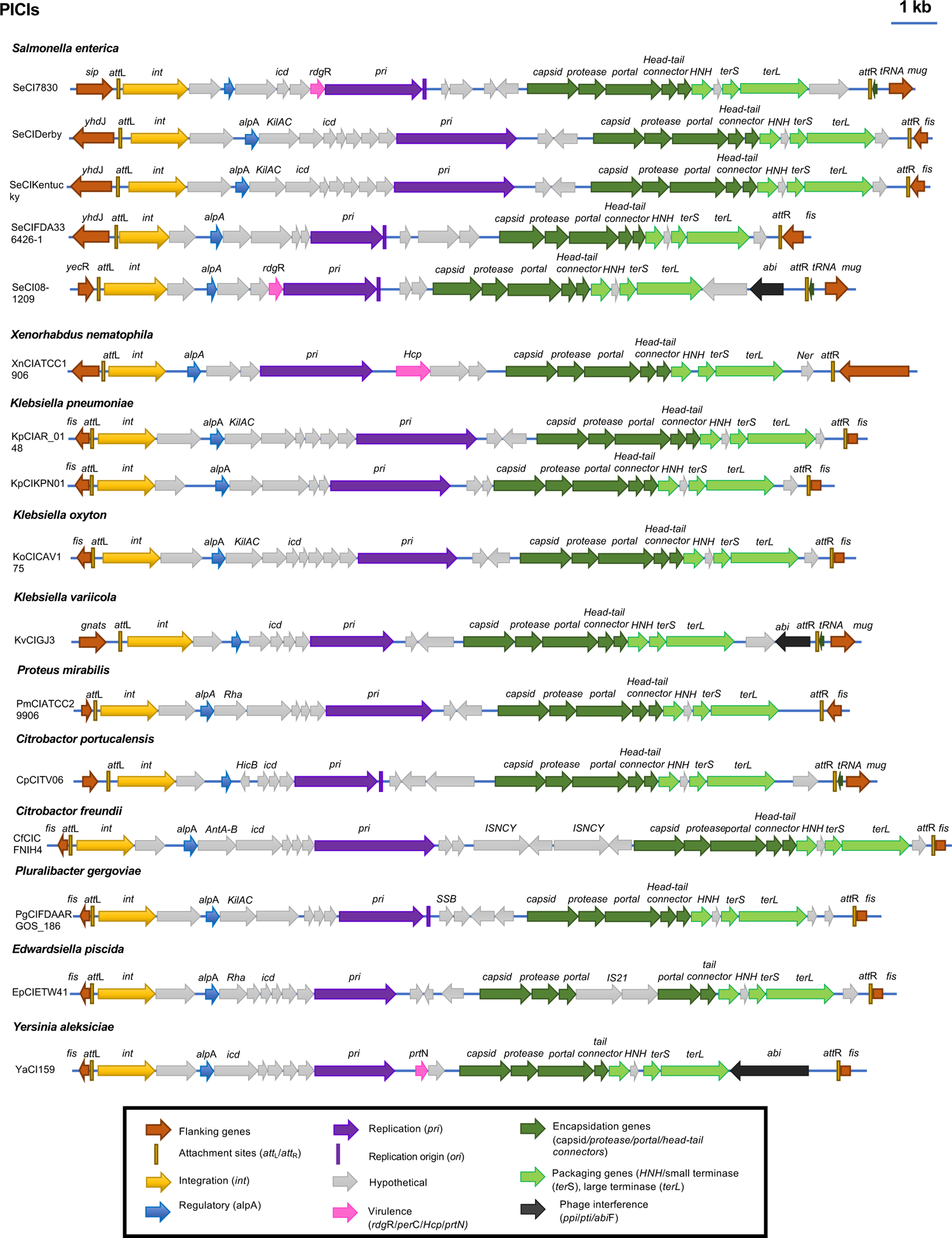
Genome maps for cf-PICIs, from different Proteobacteria species. Genomes are aligned according to the prophage convention, with the integrase genes (*int*) present at the left end of the maps. Genes are colored according to their sequence and function (see box for more details).

**Figure S3.**
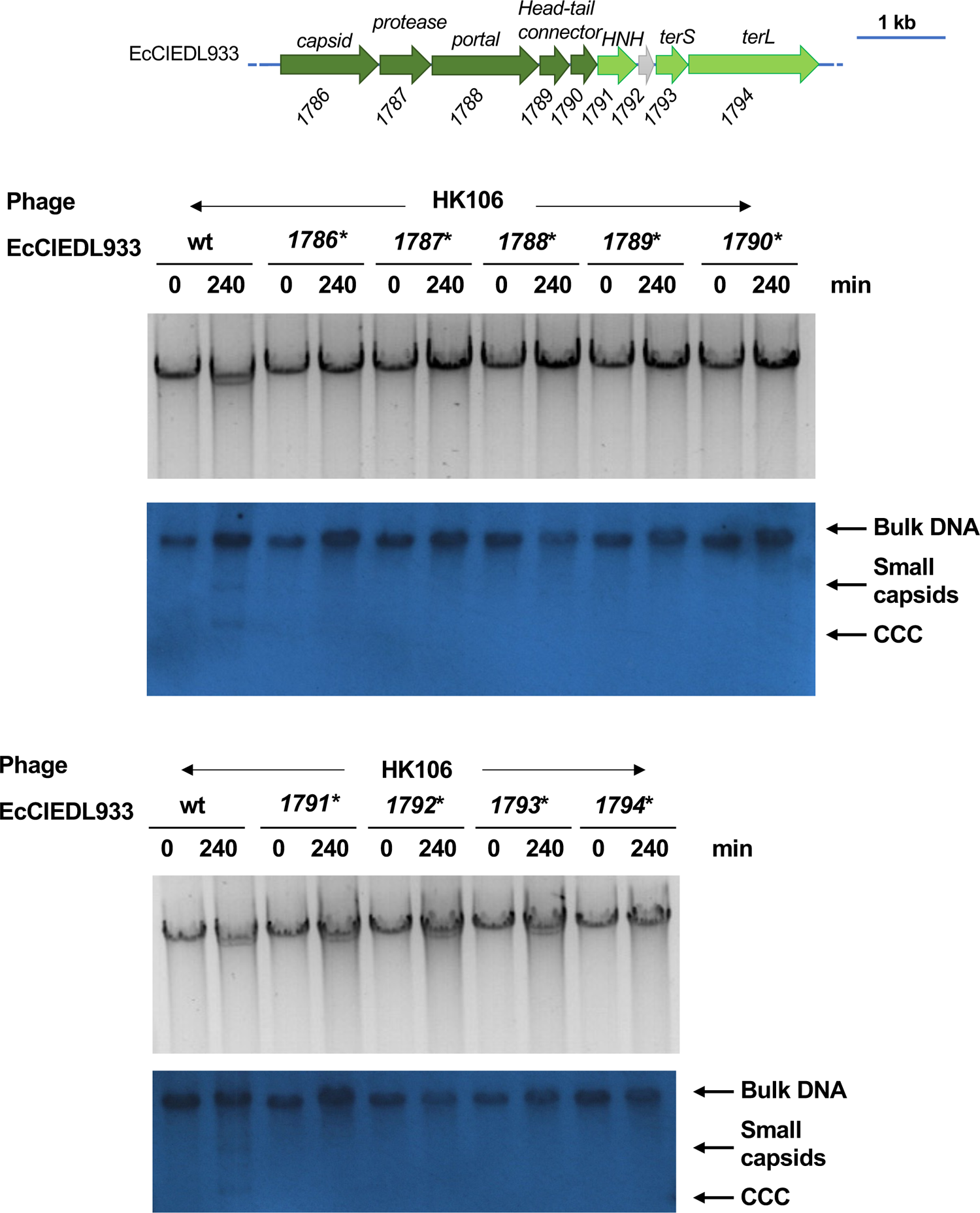
Replication of the EcCIEDL933 mutants. Lysogenic strains for phage HK106, carrying different EcCIEDL933 versions (wt or encoding individually point-mutations in the indicated genes), were MC (2 μg/ml) induced, samples were taken at the indicated time points (min) and processed to prepare minilysates, which were separated on a 0.7% agarose gel (upper panel), and Southern blotted (lower panel) with an EcCIEDL933 probe.

**Figure S4.**
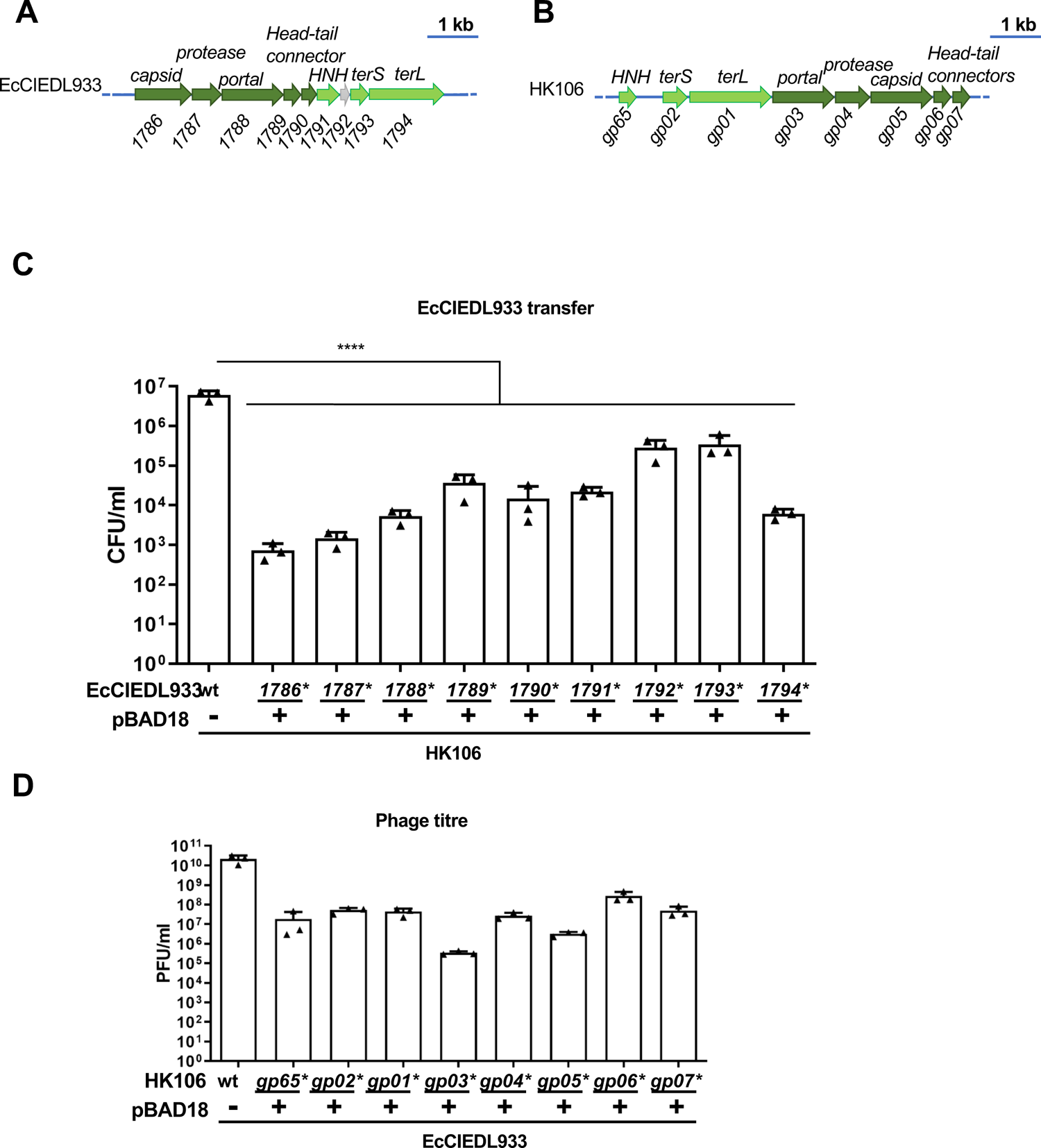
Complementation of the different EcCIEDL933 and HK106 mutants restore PICI transfer and phage titres. Genetic map of the EcCIEDL933 packaging module **(A)** or phage HK106 packaging module **(B)**. (C) Lysogenic strains for phage HK106 carrying different EcCIEDL933 versions (wt or mutants) were complemented with pBAD18 derivative plasmids expressing the different genes under study (+). Then, these strains were MC (2 µg/ml) and arabinose (0.02%) induced and the transfer of the different islands analysed using *E. coli* 594 as recipient. (D) Lysogenic strains for phage HK106 (wt or mutants), carrying wt EcCIEDL933, were complemented with the corresponding pBAD18 derivative plasmid expressing the indicated phage gene (+). The strains were MC (2 µg/ml) and arabinose (0.02%) induced and the phage titres determined using *E. coli* 594 as recipient. The means of the colony forming units (CFUs) or phage forming units (PFUs) and SD of three independent experiments are presented (n=3). A one-way ANOVA with Dunnett’s multiple comparisons test was performed to compare mean differences between wt and individually point-mutations. Adjusted p values were as follows: *ns*>0.05; *p≤0.05; **p≤0.01; ***p≤0.001; ****p≤0.0001.

**Figure S5.**
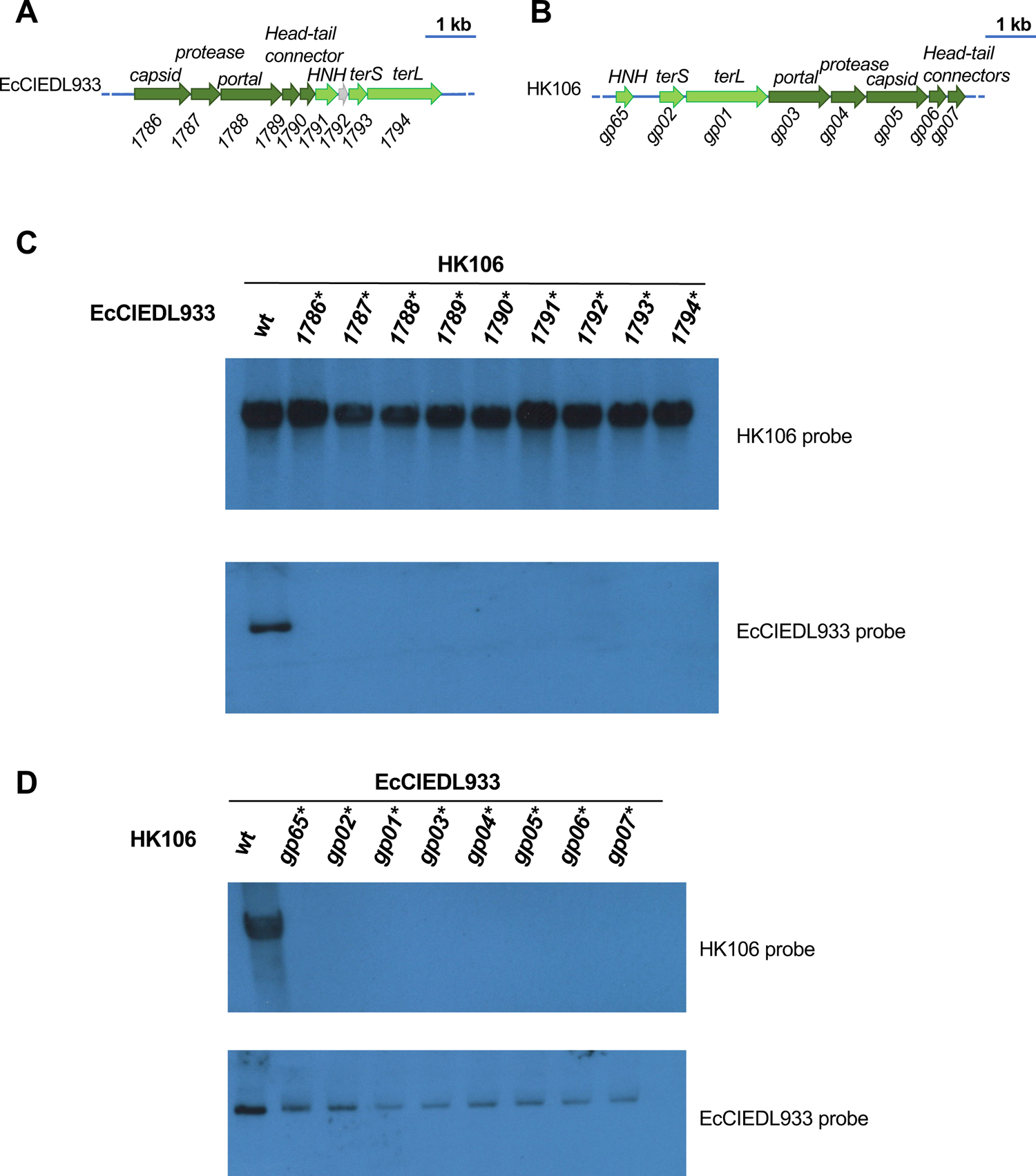
Effect of EcCIEDL933 or phage HK106 gene mutations on the formation of PICI or phage infective particles. Genetic map of the EcCIEDL933 packaging module **(A)** or phage HK106 packaging module **(B)**. **(C)** Lysogenic strains for phage HK106, carrying the wt of the different EcCIEDL933 mutant versions, were MC induced, and the DNA from the infective particles purified and analysed by Southern blot, using phage- or PICI-specific probes. **(D)** EcCIEDL933-positive strains, lysogenic for different version of the HK106 prophage (wt or carrying the indicated mutations), were MC induced, and the DNA from the infective particles purified and analysed by Southern blot, using phage- or PICI-specific probes.

**Figure S6.**
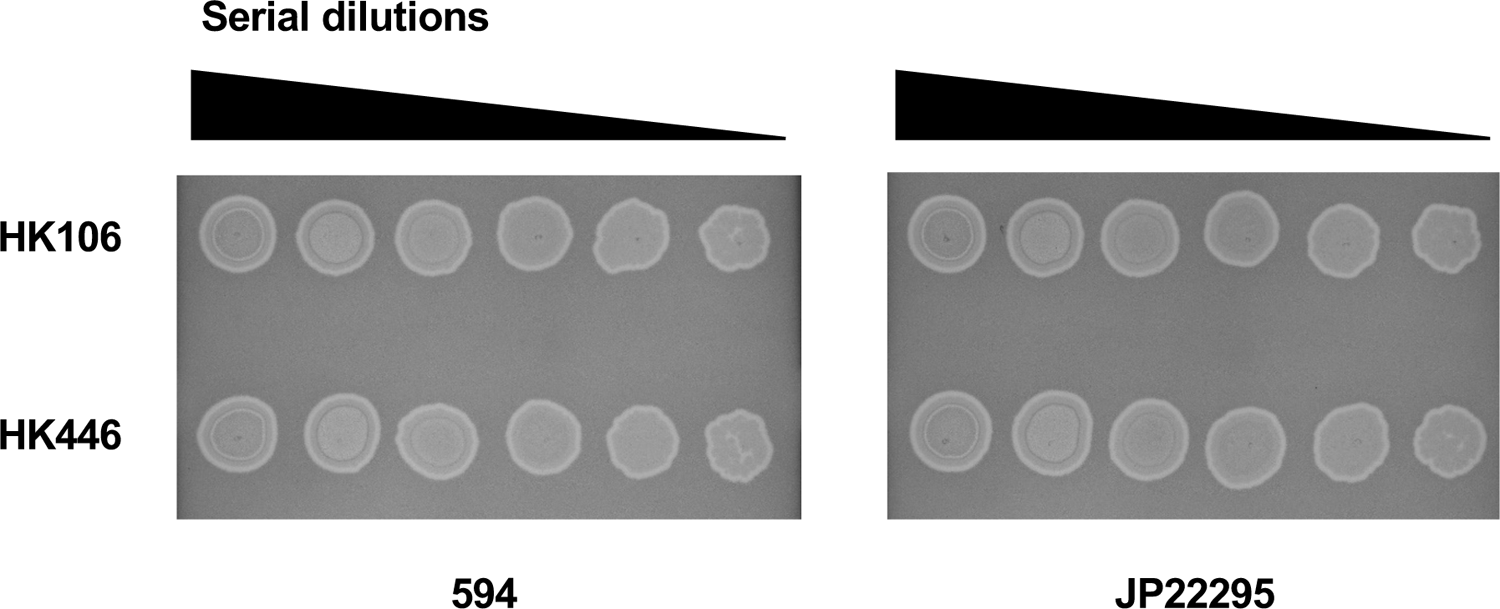
EcCIEDL933 does not block helper phage reproduction. Phage HK106 and HK446 dilutions were spotted on non-lysogenic *E. coli* strain 594, or on strain JP22295 (594 EcCIEDL933 *cat* positive).

**Figure S7.**
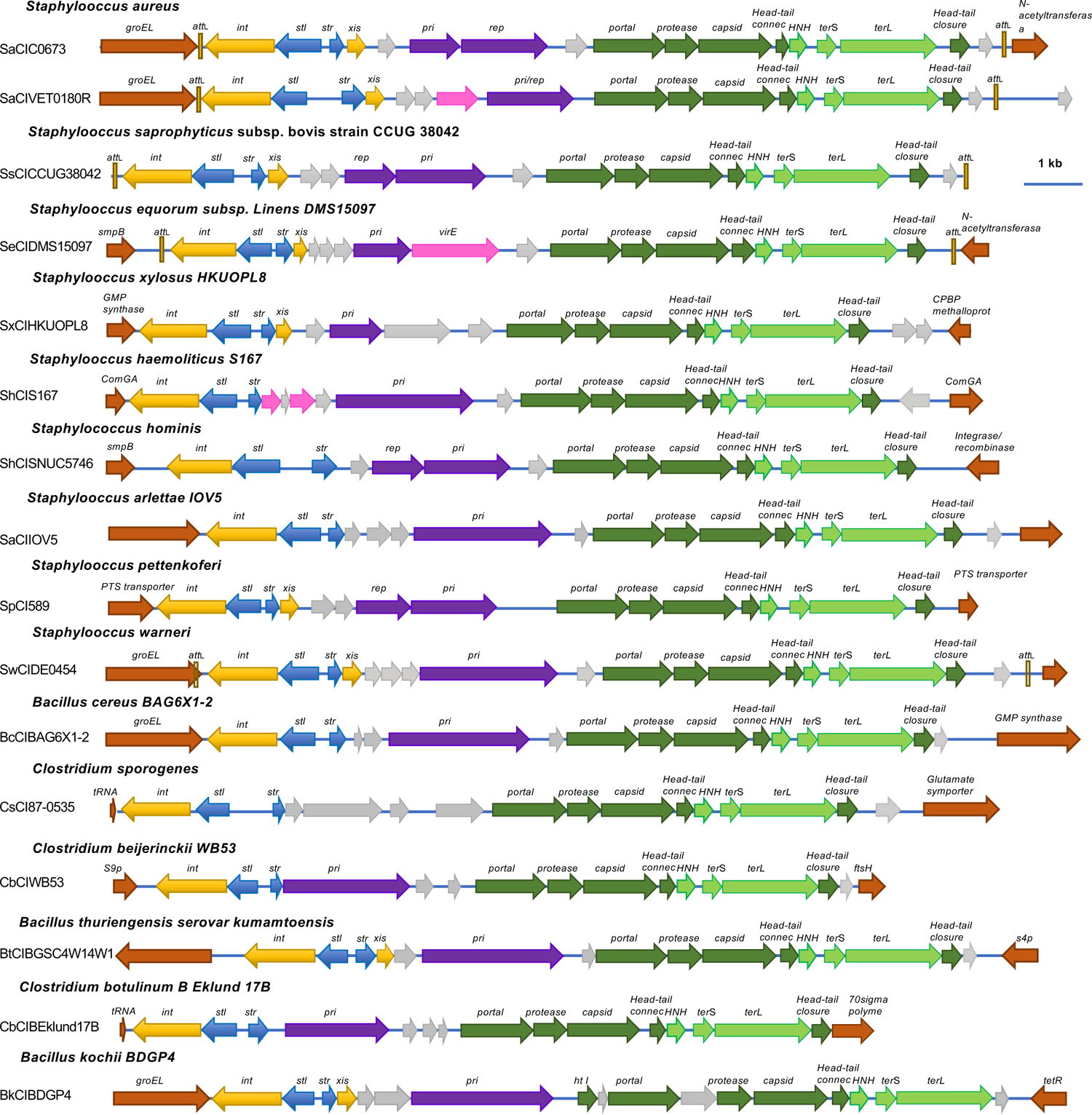
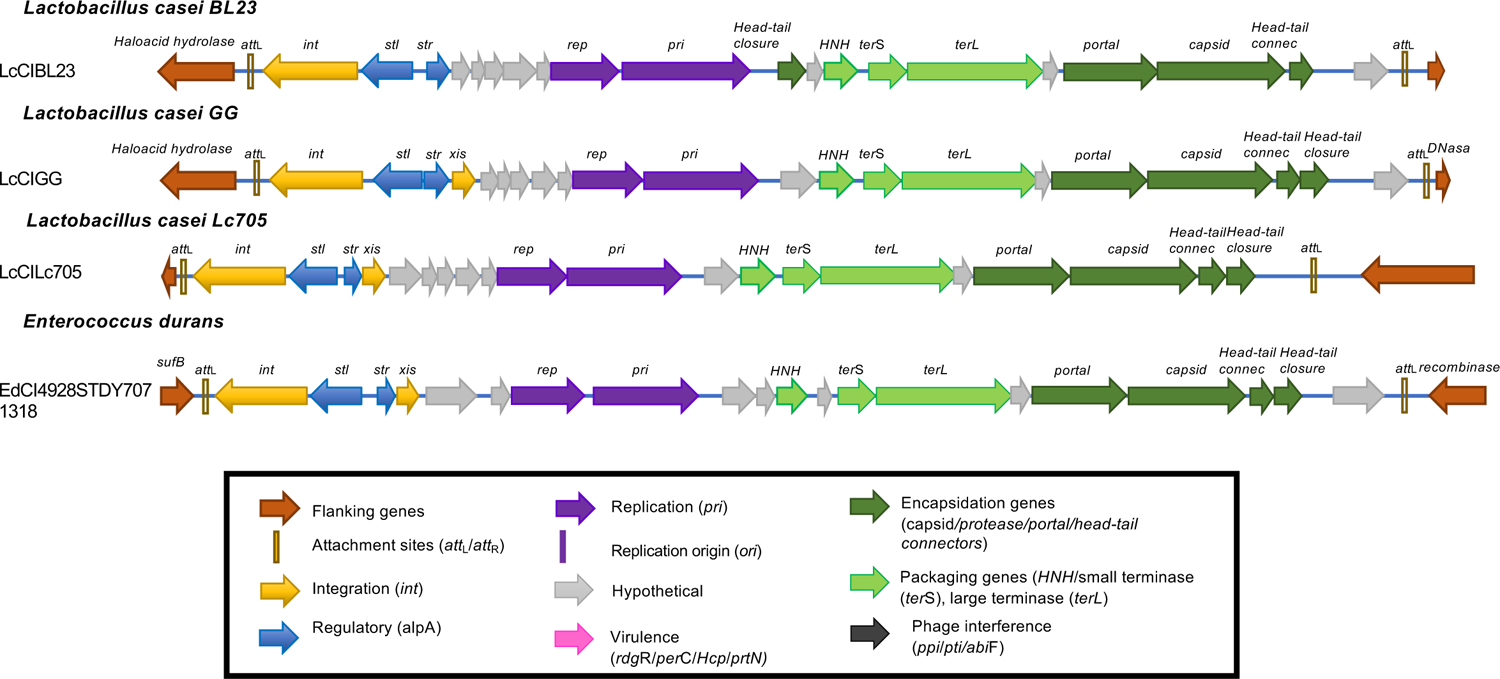
Genome maps for cf-PICIs present in Firmicutes. Genomes are aligned according to the prophage convention, with the integrase genes (*int*) present at the left end of the islands. Genes are colored according to their sequence and function (see box for more details).

**Figure S8.**
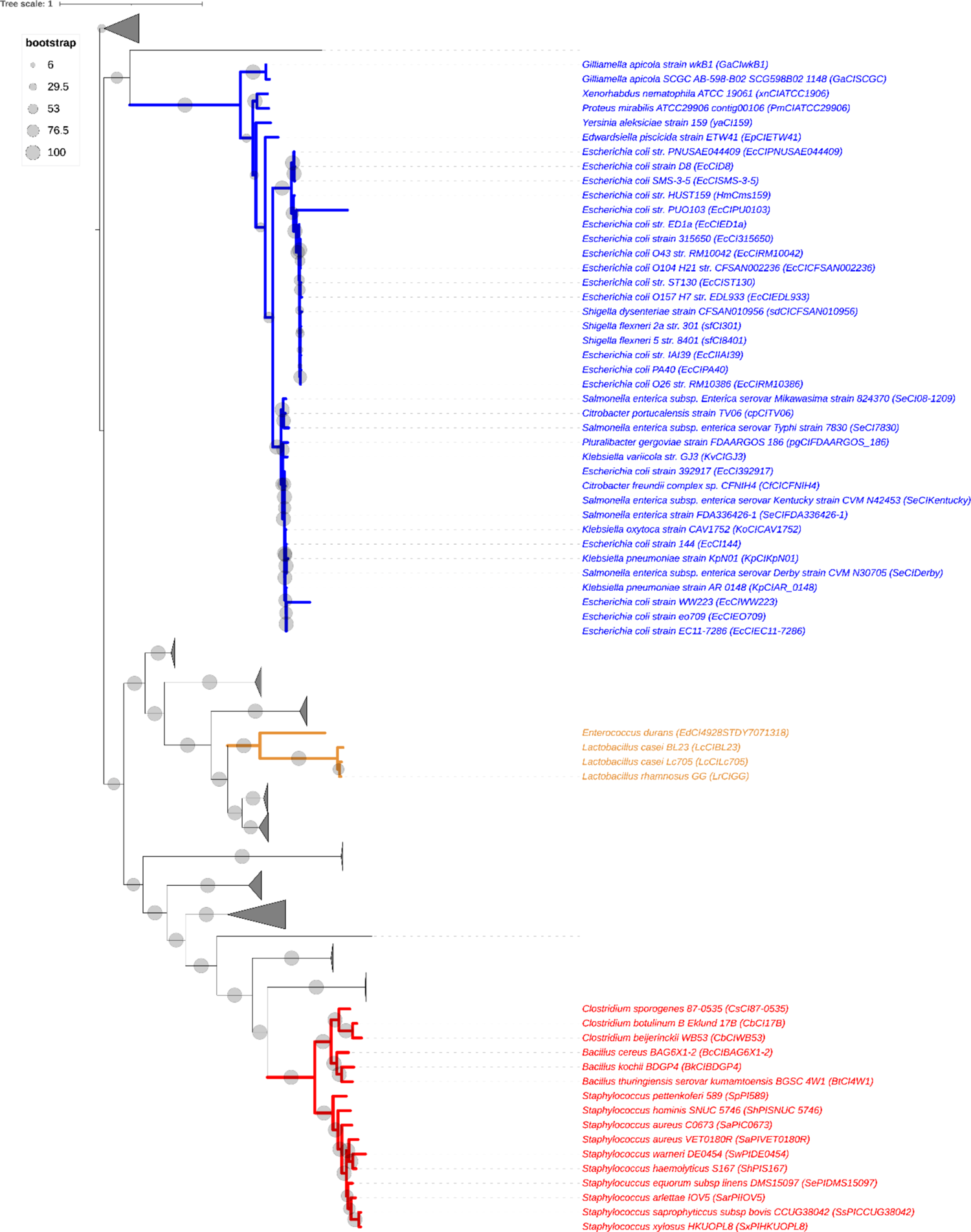
TerL of cf-PICI form three distinct phylogentic groups and are evolutionarily separated from phage homologs. Phylogenetic trees inferred from the alignment of 60 TerL homologs from cf-PICI and the best 420 phage TerL homologs. Phages branches are collapsed and shown as triangles. The different colors on the branches indicate the three different clades, formed by either Gammaproteobacteria (in blue), Lactobacillus (in orange) and a broader set of Firmicutes (in red). Bootstrap values for the main branches are indicated as grey circles. The tree was visualized and edited in iTOL ((Letunic et al., 2021).

## Notes

### Competing Interest Statement

The authors have declared no competing interest.

